# Automated analysis of whole brain vasculature using machine learning

**DOI:** 10.1101/613257

**Authors:** Mihail Ivilinov Todorov, Johannes C. Paetzold, Oliver Schoppe, Giles Tetteh, Velizar Efremov, Katalin Völgyi, Marco Düring, Martin Dichgans, Marie Piraud, Bjoern Menze, Ali Ertürk

## Abstract

Tissue clearing methods enable imaging of intact biological specimens without sectioning. However, reliable and scalable analysis of such large imaging data in 3D remains a challenge. Towards this goal, we developed a deep learning-based framework to quantify and analyze the brain vasculature, named Vessel Segmentation & Analysis Pipeline (VesSAP). Our pipeline uses a fully convolutional network with a transfer learning approach for segmentation. We systematically analyzed vascular features of the whole brains including their length, bifurcation points and radius at the micrometer scale by registering them to the Allen mouse brain atlas. We reported the first evidence of secondary intracranial collateral vascularization in CD1-Elite mice and found reduced vascularization in the brainstem as compared to the cerebrum. VesSAP thus enables unbiased and scalable quantifications for the angioarchitecture of the cleared intact mouse brain and yields new biological insights related to the vascular brain function.

**GRAPHICAL ABSTRACT:** **Supporting material of VesSAP is available at http://DISCOtechnologies.org/VesSAP**

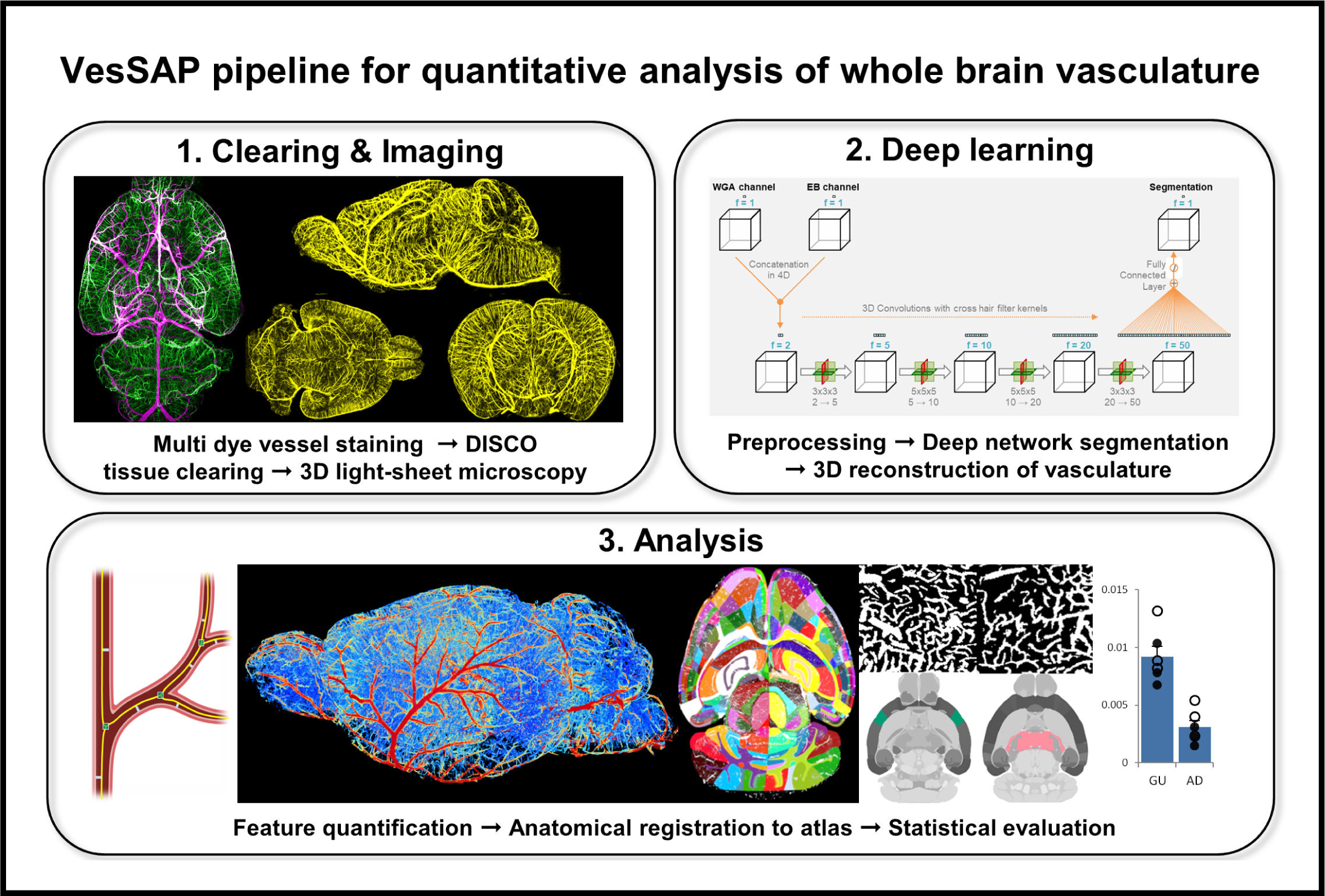

## INTRODUCTION

Changes in the brain vasculature are a key feature of a large number of diseases effecting the brain. Primary angiopathies, vascular risk factors (e.g., diabetes), traumatic brain injury, vascular occlusion and stroke all affect the brain vascular network and interfere with normal microcirculation and vascular function^1-5^. Alterations of the brain microvasculature are also seen in neurodegenerative diseases, such as Alzheimer’s disease, tauopathy and amyloidopathy. These hallmarks of the Alzheimer’s disease, can lead to aberrant remodeling of the blood vessels^1,6-8^. Consequently, capillary rarefaction is frequently used as a marker for vascular damages^9^. Thus, quantitative analysis of the entire brain vasculature including the capillary bed is pivotal to develop a better understanding of physiological and pathological brain function. However, quantifying micrometer scale changes in the cerebrovascular network of intact brains has been difficult for two main reasons.

First, labeling and imaging of the complete mouse brain vasculature down to the smallest blood vessels has to be achieved. Magnetic resonance imaging (MRI), for instance, does not have sufficient resolution to capture capillaries^10^. MicroCT imaging can visualize the microvasculature, but because of specimen size constraints it fails to acquire a whole intact mouse brain^11^. Fluorescent microscopy, on the other hand, provides a higher resolution but can typically be applied to 1-200 μm thin tissue slices, which does not pre-serve the structure of an end-to-end vascular network. Recent advances in tissue clearing could overcome this problem, but so far there has been no demonstration of all vessels of all sizes in an entire brain in three dimensions (3D)^12^.

The second challenge relates to automated analysis of 3D imaging data for structures that are spanning entire mouse brains, which cannot be analyzed piece by piece in a reliable and scalable manner. Scanning transparent specimens of several millimeters size at micrometer resolution, inevitably introduces substantial variance in the signal intensity and signal-to-noise ratio at different depths. Thresholding methods are not capable of segmenting these large scans, whereas shape-based filtering approaches such as Frangi filters cannot reliably identify vessels from background^13,14^. To overcome these limitations more advanced image processing methods with local spatial regularization have been proposed for processing light-sheet scans^15^. However, such methods including local spatial regularization cannot segment large vascular networks across changing intensity distributions. Finally, the size of the acquired datasets poses a difficulty to assess the organization of the whole vascular network; therefore, such methods can only segment small volumes^15-19^.

Here, we present VesSAP (**Ves**sel **S**egmentation & **A**nalysis **P**ipeline), a method for automated quantitative analysis of the entire mouse brain vasculature, which overcomes the limitations stated above. To achieve this, we first developed a dual vascular staining approach using wheat germ agglutinin (WGA) and Evans blue (EB) to stain both small and large vessels in two fluorescent channels, consistently throughout the entire brain. Next, we cleared whole stained brains using the 3DISCO method^20^ and imaged them with light-sheet microscopy at micrometer resolution. Second, we developed a deep fully convolutional network (FCN), which exploits the imaging data from both dyes to provide a high-quality segmentation of the vasculature in 3D. Subsequent feature extraction and registration to the latest Allen adult mouse brain atlas enabled us to quantify all features of interest with respect to their topographical location. Our deep learning-based approach works reliably despite variations in signal intensities, outperforming previous filter-based methods and reaching the quality of segmentations of human annotators. To our knowledge, this is the first time that a deep learning approach is being used to analyze complex imaging data of cleared mouse brains i.e. spanning the entire brain end-to-end.

We further applied VesSAP to a set of 6 mice from two commonly used mouse strains to systematically explore strain-related differences in vascular anatomy across brain regions as described by the Allen brain atlas. We reported new biological findings and provide a comprehensive reference set of vessel anatomy features, revealing unique structures of different brain regions. Thus, VesSAP represents an integrated pipeline enabling automated and scalable analysis of the complete mouse brain vasculature (**Fig. 1**). All parts of the VesSAP are publicly hosted online for easy adoption, including the imaging protocol, the data (original scans, registered atlas data), the trained algorithms, training data and a reference set of features describing the vascular network in all brain regions at the following address: http://DISCOtechnologies.org/VesSAP

**Figure 1:**
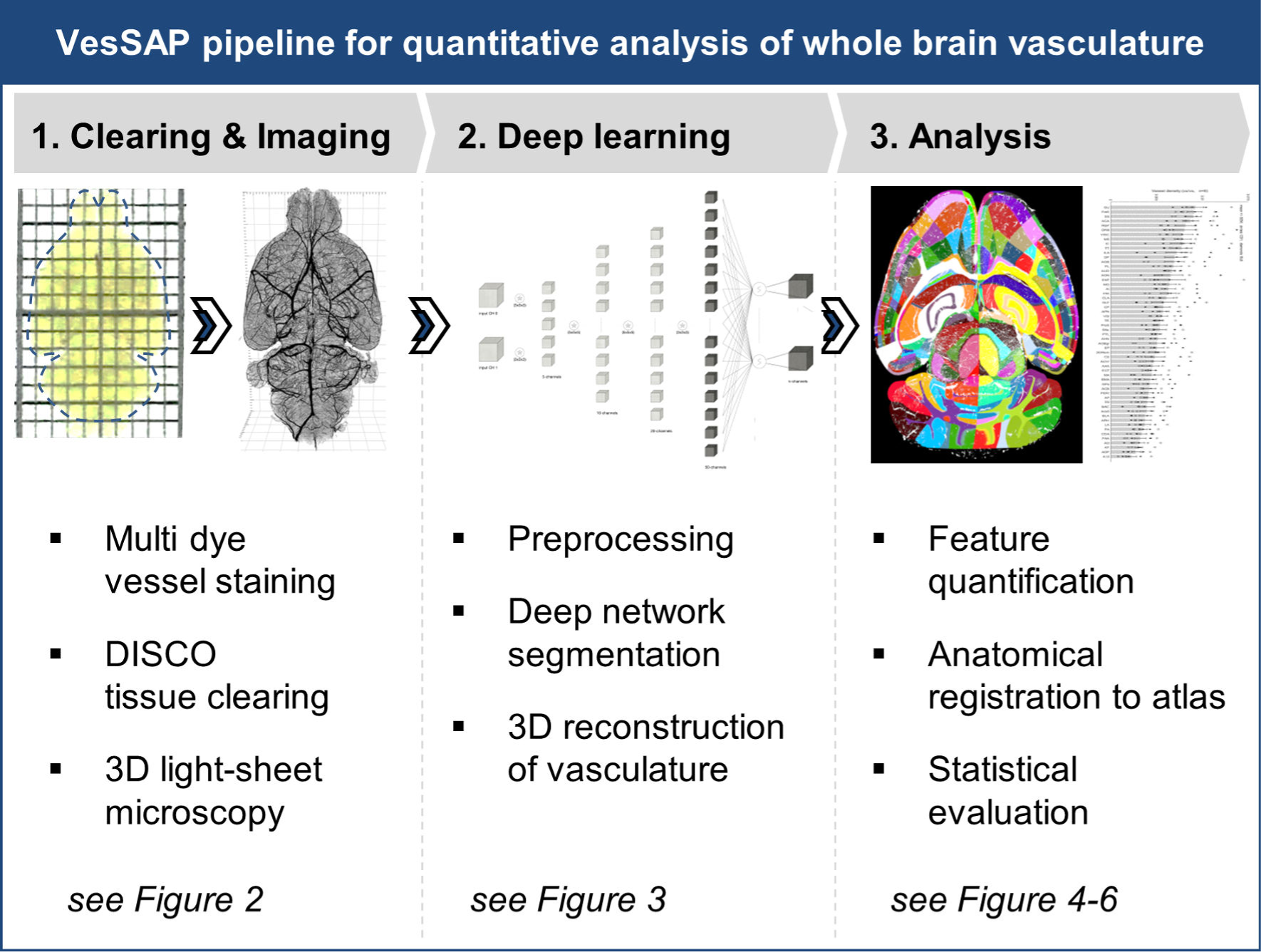
Summary of the VesSAP pipeline for automated whole brain analysis of the perfused vasculature. The proposed method consist of three modular steps: **1,** multi dye vessel staining and DISCO tissue clearing for high imaging quality using 3D light-sheet microscopy; **2,** Deep-learning based segmentation of blood vessels with 3D reconstruction and **3**, Anatomical feature extraction and mapping of the entire vasculature to the Allen adult mouse brain atlas for statistical analysis.

## RESULTS

Tissue clearing methods enable imaging of un-sectioned biological specimens. To extract biologically meaningful data, they have to be combined with reliable and automated image analysis methods. Towards this goal, we developed VesSAP, a deep learning-based method to accurately and automatically analyze the vasculature of cleared mouse brains. VesSAP encompasses 3 major steps: 1) staining, clearing and imaging of the mouse brain vasculature by two different dyes (WGA and EB) down to the capillary level, 2) transfer learning-based algorithms to automatically segment and trace the whole brain vasculature data at the capillary level and 3) extraction of vascular features for hundreds of brain regions by registering the data to the Allen brain atlas (**Fig. 1**). We applied VesSAP to generate vascular reference maps for two commonly used mouse strains under physiological conditions: C57BL/6J and CD1-Elite mice. We report the first evidence of secondary intracranial collateral vascularization in CD1-Elite mice. Furthermore, our work shows a significantly decreased vascularization density in the brainstem as compared to the cerebrum in both mouse strains.

### Step 1: Vascular staining, DISCO clearing, and imaging

Towards staining the entire vasculature, we applied a combination of two dyes, WGA and EB staining in two fluorescent channels. We then performed 3DISCO clearing^21^ and light-sheet microscopy imaging of whole mouse brains at micrometer resolution (**Fig. 2A-C, Supporting Fig. 1**). WGA predominantly stains small vessels and, importantly, captures even the smallest ca-pillaries down to diameters of a few micrometers, while EB predominantly stains large vessels including the middle cerebral artery and the circle of Willis (**Fig. 2D-I, Supporting Fig. 2**). Merging the signals from both dyes yields a staining of the complete vasculature, showing the complementary nature of both dyes (**Fig. 2C,F**). Importantly, the signals from both dyes are perfectly congruent when staining the same vessel and solely come from the vessel wall layer (**Fig. 2G-I, Supporting Fig. 2**). Furthermore, owing to the dual labeling, we maximized the signal to noise ratio (SNR) for each dye independently to avoid saturation of differently sized vessels when only a single channel is used. We achieved this by independently optimizing the excitation and emission power. For WGA, we reached a higher SNR for small capillaries; bigger vessels, however, were barely visible (**Supporting Fig. 3**). For EB, the SNR for small capillaries was substantially lower but larger vessels reached a high SNR (**Supporting Fig. 3**). Thus, integrating the infor-mation from both channels allows homogenous staining of the entire vasculature throughout the whole brain, and results in a high SNR for high-quality segmentations and analysis.

**Figure 2:**
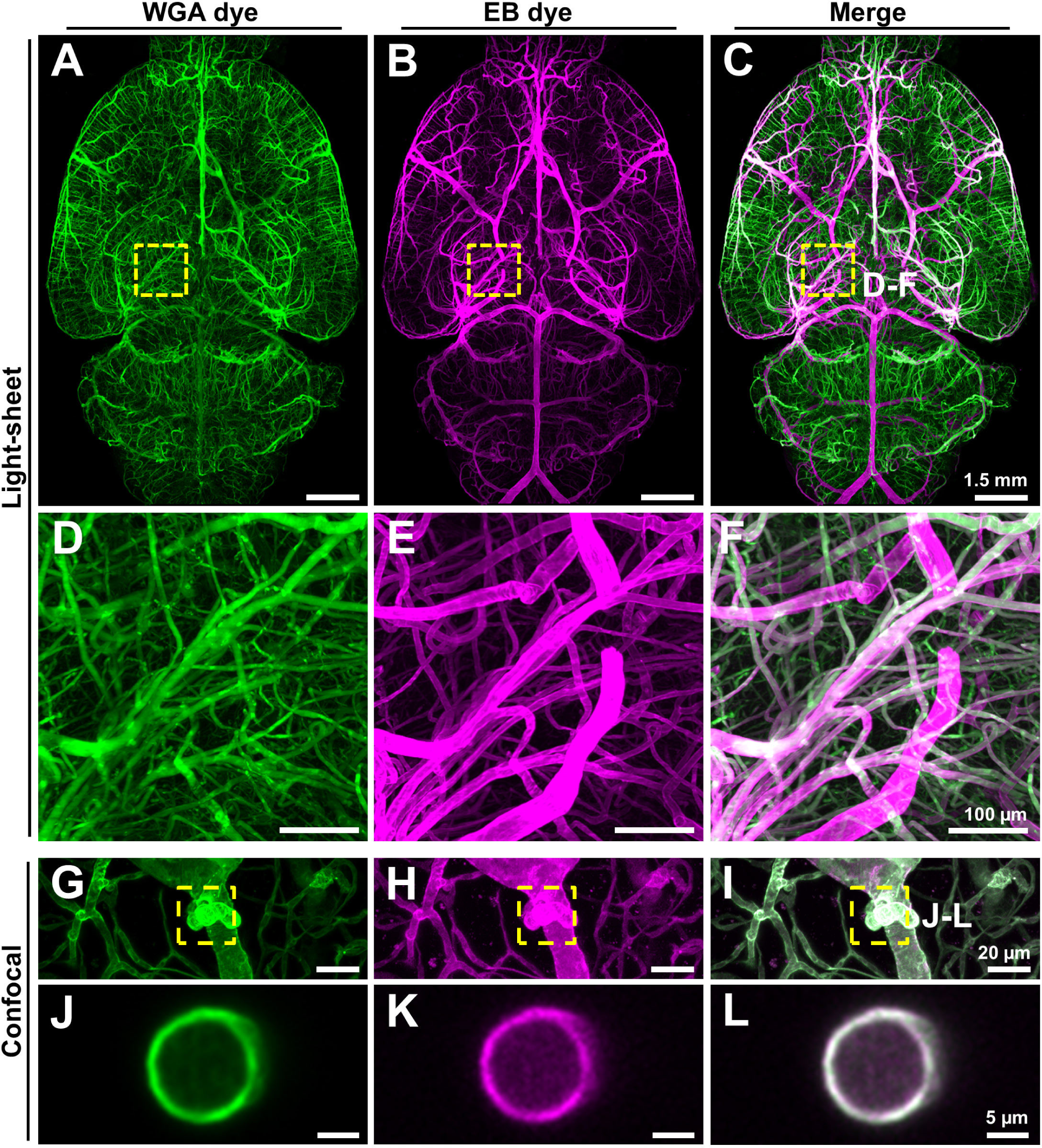
Enhancement of vascular staining using two complementary dyes. **A-C**, Maximum intensity projections of the automatically reconstructed tiling scans of WGA (**A**) and Evans blue (**B**) signal in the same sample reveal all details of the perfused vascular network in the merged view (**C**). D-F: Zoom-ins from marked region in (**C**) showing fine details. **G-L**, Confocal microscopy confirms that WGA and EB dyes stain the vascular wall (**G-I**, maximum intensity projections of 112 µm) and that the vessels retain their tubular shape (**J-L**, single slice of 1 µm).

### Step 2: Segmentation of the volumetric images

To enable extraction of quantitative features of the vascular structure, the vessels in the acquired brain scans need to be segmented in 3D. Motivated by the recent success of deep learning based approaches in biomedical image data analysis^22-32^, including human MRI data segmentation, we developed a deep FCN to exploit the complementary signals of both dyes to derive a complete segmentation of the entire vasculature.

Our network architecture is a deep FCN, which segments vessels based on the input from two imaging channels. The network consists of 5 layers, 4 convolutional layers followed by one fully connected layer. In a first step, the two input channels (WGA and EB) are concatenated. This yields a set in which each voxel in 3D space is characterized by two features. Each convolutional step integrates the information from the voxel’s 3D neighborhood. Here we use cross-hair shaped convolutional kernels to sample the local surrounding of each voxel in a sparse manner along three orthogonal planes^25^ (**Fig. 3A**). After the last convolution, the information from 50 features per voxel is combined with a fully connected layer and a sigmoidal activation to estimate the likelihood that a given voxel represents a vessel. Subsequent binarization yields the final segmentation. In both, training and testing, the images are processed on sub-patches of 50 × 100 × 100 pixels.

**Figure 3:**
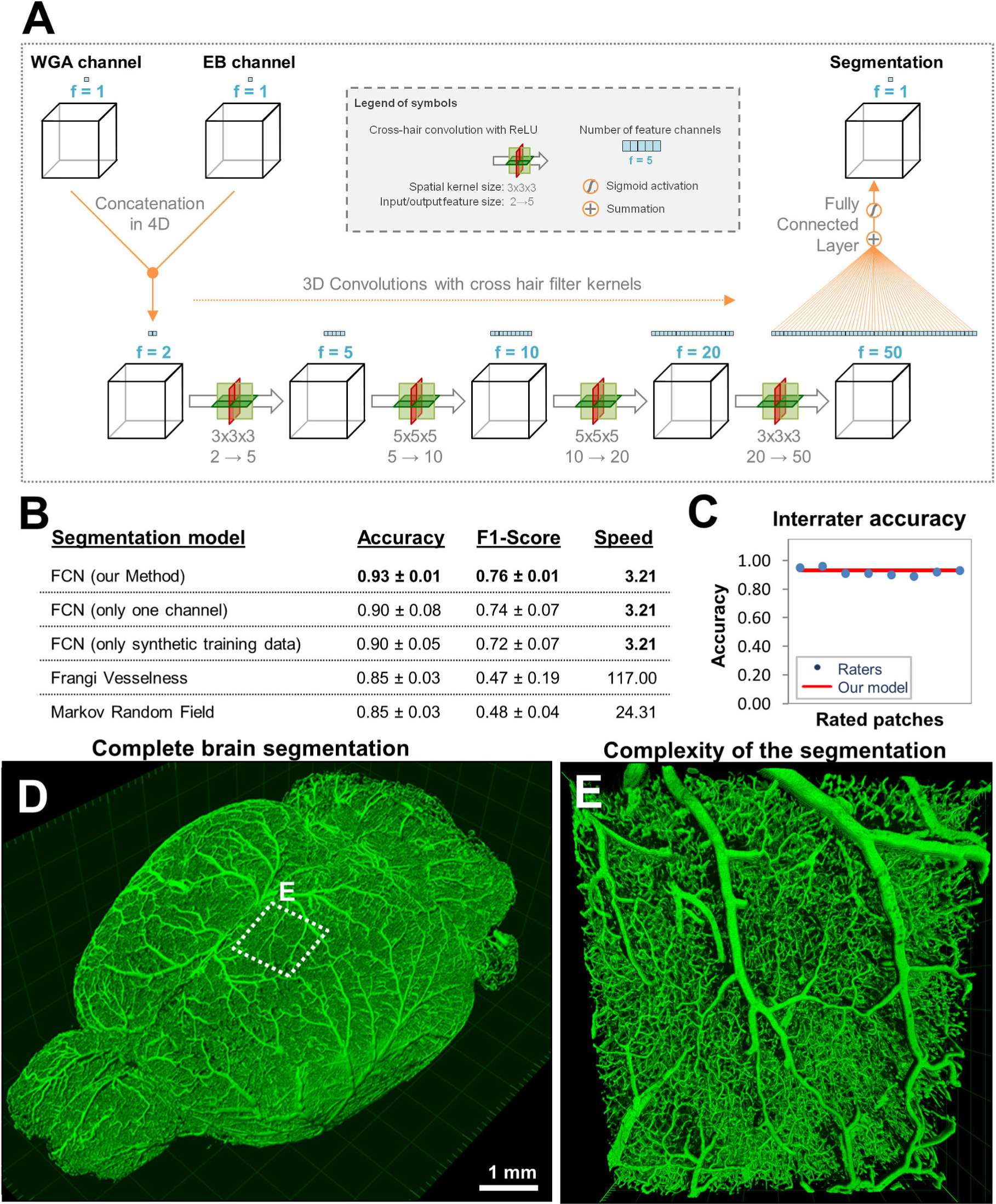
Deep learning architecture of VesSAP and performance on vessel segmentation. **A**, The proposed VesSAP network architecture, consisting of four convolutional layers and a sigmoid classification layer. Schematic representation of the convolutional cross hair operations applied in each convolutional step. **B**, Evaluation metrics for the performance of VesSAP compared to a one channel architecture and the commonly used Frangi filter and speed of segmentation for one image volume of 500 × 500 × 50 pixels. **C**, The quantification for interrater experiment showing the performance of our model compared to four human experts. Each of them annotated two patches. **D**, 3D rendering of full brain segmentation from a CD1-E mouse. E, 3D rendering of a small patch marked in (**D**) showing large vessels and connected capillaries.

Often, deep neural networks require large amount of annotated data to be trained. Here, we circumvented this requirement with a transfer learning approach, which is frequently used to train algorithms with limited annotated data^33^. In short, the network was first pre-trained on a large, synthetically generated vessel-like data set (**Supporting Fig. 4**)^34^ and then refined on a small amount of manually annotated part of the real brain vessel scans. This approach drastically reduced the need for manually annotated training data, as indicated by the following observations: first, the network that was solely trained on synthetic data already yields acceptable segmentations of real brain vessel scans. Second, the refinement on the real data set already converged after a few training epochs. Third, only a very small amount of manually annotated training data (here: 0.02 % of a full brain scan) was needed to segment the vasculature of an entire brain with the quality of human annotations.

To assess the quality of the segmentation quantitatively, we compared the network prediction with manually created ground truth segmentation and ran several experiments with alternative approaches (**Fig. 3B, Supporting Fig. 5**). To provide comparability with the literature, we reported two voxel-wise measures that quantify the quality of the segmentation, accuracy and the F1-score^35^. As compared with the ground truth, our network exploiting the information from WGA and EB achieved an accuracy of 0.93 ± 0.01 and a F1-Score 0.76 ± 0.01 (**Fig. 3B**). To assess the importance of integrating information from both, WGA and EB, we designed a control network that only has access to the commonly used WGA signal, which reduced the quality of the vessel segmentation (accuracy: 0.90 ± 0.08; F1-Score 0.74 ± 0.07). As further controls, we implemented alternative state-of-the-art methods and found that our network outperforms classical Frangi filters^13^ (accuracy: 0.85 ± 0.03; F1-Score 0.47 ± 0.18), as well as recent methods considering local spatial context via Markov random fields^15,36^ (accuracy: 0.85 ± 0.03; F1-Score 0.48 ± 0.04).

Next, we compared the segmentation accuracy of our network with human annotations. A total of 4 human experts independently annotated two volumes each and these segmentations were again compared to the ground-truth segmentation. We found that the accuracy of segmentation quality of independent human annotators was comparable to the predicted segmentation of our network (**Fig. 3c**). Moreover, our network could segment a whole mouse brain scan within 24 hours on a single GPU workstation whereas the annotators who created our segmentations would need more than a year to process a whole brain. In summary, the VesSAP pipeline is able to segment the whole brains vasculature at human level accuracy with a substantially higher speed.

**Fig. 3D** and **Supporting Video 1,2** show an example of a brain vasculature that was segmented by VesSAP in 3D. Zooming into a smaller patch reveals that the connectivity of the vascular network was fully maintained (**Fig. 3E, Supporting Video 1**). Comparing single slices of the imaging data with the predicted segmentation shows that vessels are accurately segmented regardless of absolute illumination and vessel diameter (**Supporting Fig. 6**).

### Step 3: Feature extraction and atlas registration

It has previously been shown that vessel density, radius and the number of bifurcation points can be used to describe vascular anatomy^3^. Hence, we used our segmentation to trace and quantify these features as the distinct parameters to characterize the mouse brain vasculature (**Fig. 4A, Supporting Video 3**). A visualization of the quantified vessel radius along the entire vascular network is shown in **Fig. 4B**. After extracting vascular features of the whole brain with VesSAP, we registered the volume to the Allen brain atlas (**Supporting Video 4,5**). This allowed us to map the segmented vasculature and corresponding features to distinct anatomical brain regions. A representative cross-section of the brain through the vasculature, color-coded by coarse anatomical regions, is depicted in **Fig. 4C.** Each anatomical region can be further divided into subregions, yielding a total of 1238 anatomical structures (619 per hemisphere) for the entire brain (**Fig. 4D**). This allows analyzing each denoted brain region and grouping regions into clusters such as left vs. right hemisphere, gray vs. white matter or hierarchical clusters of the Allen brain atlas ontology. For our subsequent statistical feature analysis, we chose to group the labeled structures according to the main 55 anatomical clusters of the current Allen brain atlas ontology. We thus provide the first whole mouse brain vascular map with the extracted centerlines, bifurcation points and radius down to the capillary level.

**Figure 4:**
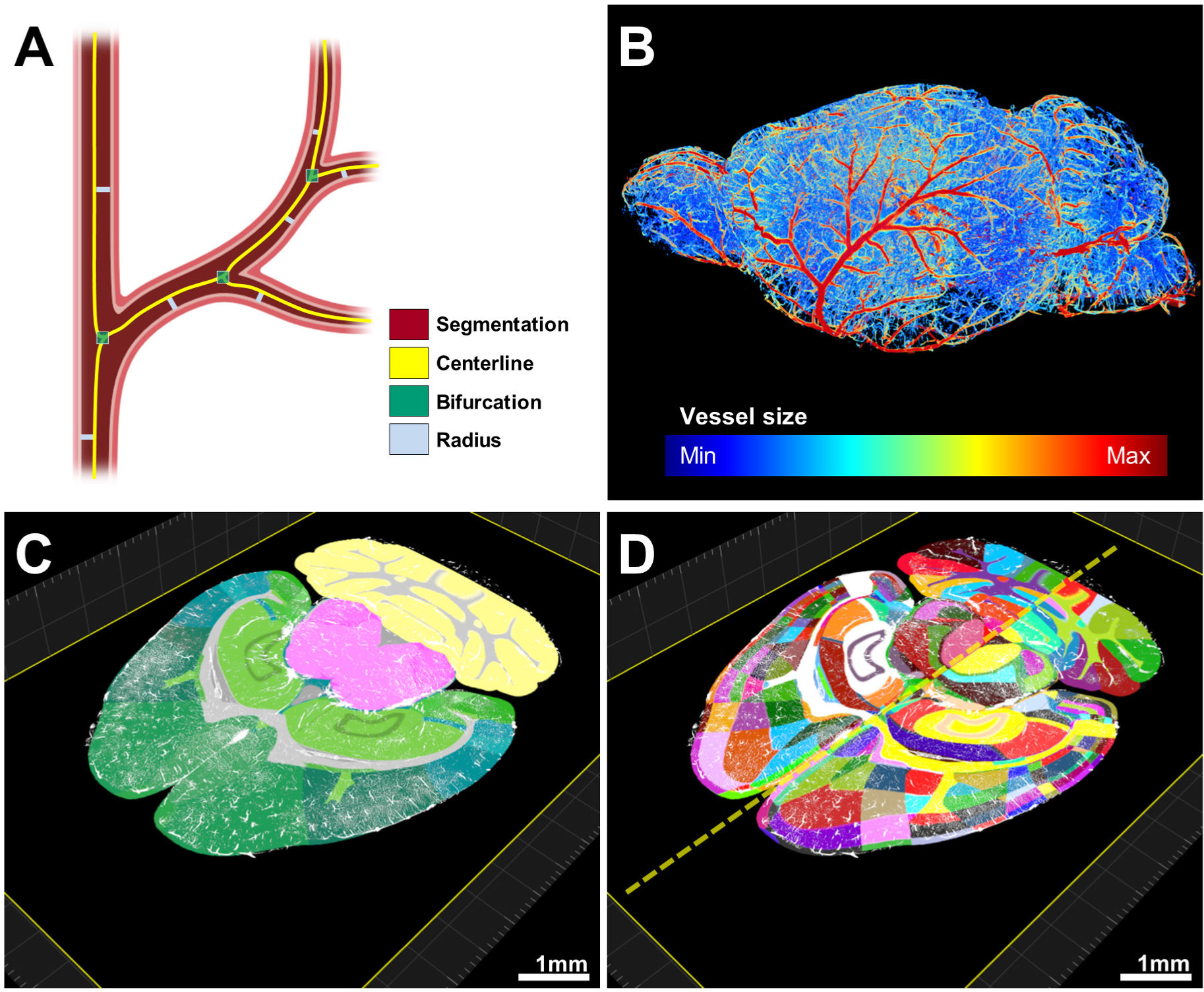
Pipeline showing feature extraction and registration process into the native space of each scan. **A**, Representation of the features extracted from vessels: volume mask, centerline, bifurcations and radii. **B**, Radius illustration of the vasculature in a representative CD1-E mouse brain. C-D, Vascular segmentation results overlaid on the hierarchically (**C**) and randomly color coded atlas to reveal all annotated regions (**D**) available including hemispheric difference (dashed line in D).

### VesSAP provides a reference map of the whole brain vasculature in mice

To extend utility of our new method, we next derived three secondary features from the segmentation to build a reference map of the brain vasculature: 1) To approximate the total length of all vessels in a given volume of interest, we extract the centerlines and count the voxels along it (expressed in voxel / voxel ratio); 2) density of bifurcation points normalized to the region size (expressed in count / mm^3^); and 3) average radius for each region (expressed in μm). These features were all referenced to the Allen brain atlas ontology. We used these features to determine the vascular features in 6 individual brain samples from the C57BL/6J and CD1-Elite strains (n = 3 mice for each strain). From these quantifications we derived the following conclusions: first, the vessel radius is evenly distributed in different regions of the same brain (**Fig. 5A**). Second, the bifurcation density and vessel length are unevenly distributed in the same brain over different regions, while they correlate well among different mice for the same regions (**Fig. 5B,C**). Moreover, bifurcation density correlates well with the vessel length ratio in most of the brain regions (**Fig. 5D**, Pearson’s correlation, *r*=0.956, p-value=1.971^-30^). Third, we observed that the extracted features show no significant statistical difference for the same anatomical cluster between the two strains (C57BL/6J and CD1-Elite) (**Supporting Fig. 7, Supporting Tables 1-4**).

**Figure 5:**
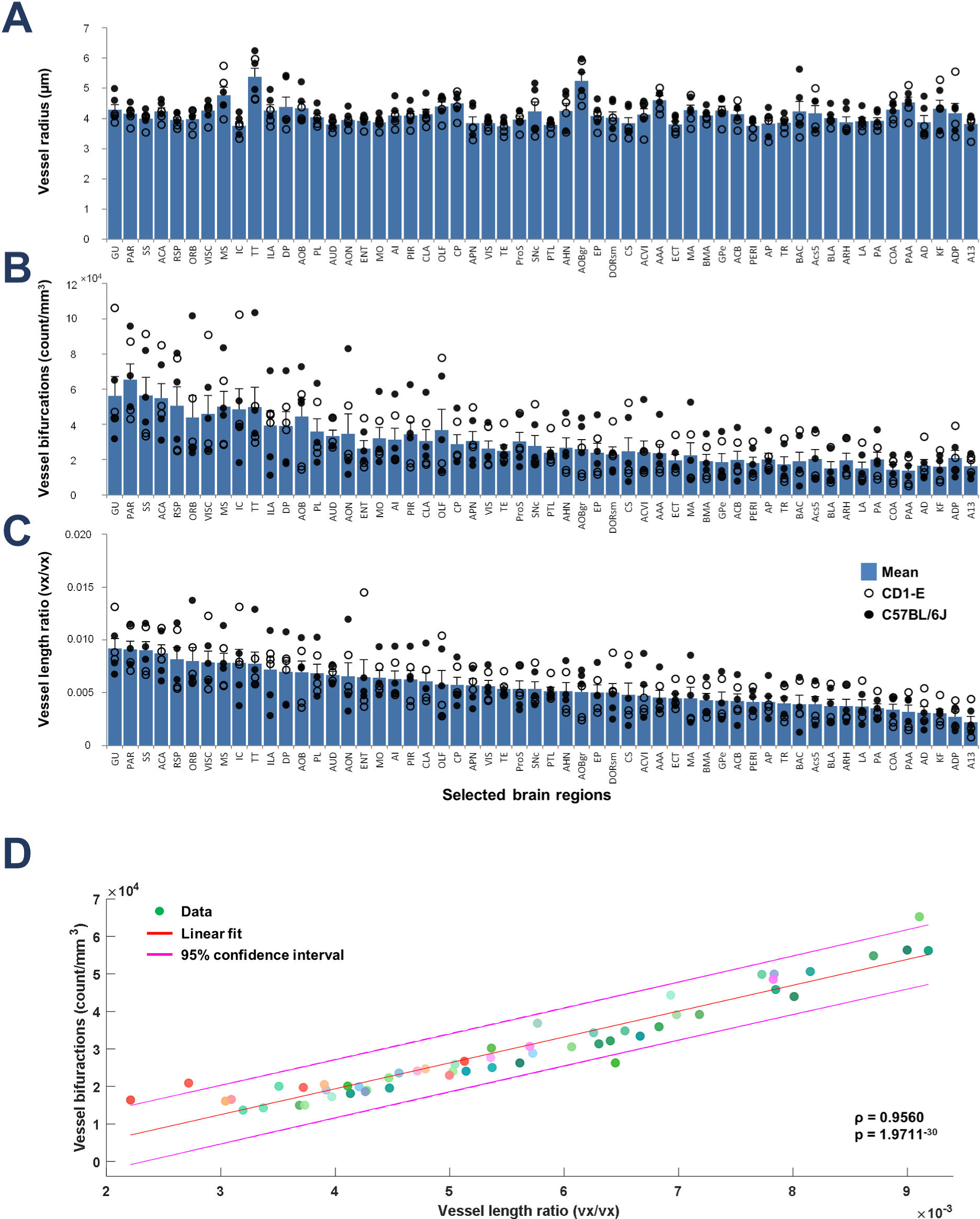
Reference values of vessel radii, bifurcations and vessel lengths in cleared tissue mapped to 55 anatomical clusters of the Allen brain atlas. **A-C**, Representation of the average radius (**A**), number of bifurcation points (**B**) and vessel length (**C**) in each of the selected anatomical clusters in the Allen brain atlas. Open circles denotes CD1-E and closed circles C57BL/6J strain; each circle represents a single mouse. As the features are similar between two strains, we pooled the data of them to generate reginal feature graphs. **D**, Scatter plot of the vessel length against the bifurcations shows a region specific tight correlation between these features (Pearson’s *r* = 0.9560; p = 1.9711-30). Each color represents a different brain area. All abbreviations are listed in the Supporting table 1.

Next, we visually inspected exemplary regions and validated the output of VesSAP. For example, the Gustatory areas showed higher vascular density compared to the Anterodorsal nucleus (**Fig. 6A**) as predicted by VesSAP (**Fig. 6B,C**). This inspection also suggested that the capillary density was the primary reason for the regional variations in the same brain. Additionally we found 1) direct intracranial vascular anastomosis in both strains (white arrowheads, **Fig. 6D,E**), and 2) that the anterior cerebral artery, middle cerebral artery and the posterior cerebral artery are connected at the dorsal visual cortex (red arrowheads, **Fig. 6D,E**).

**Figure 6:**
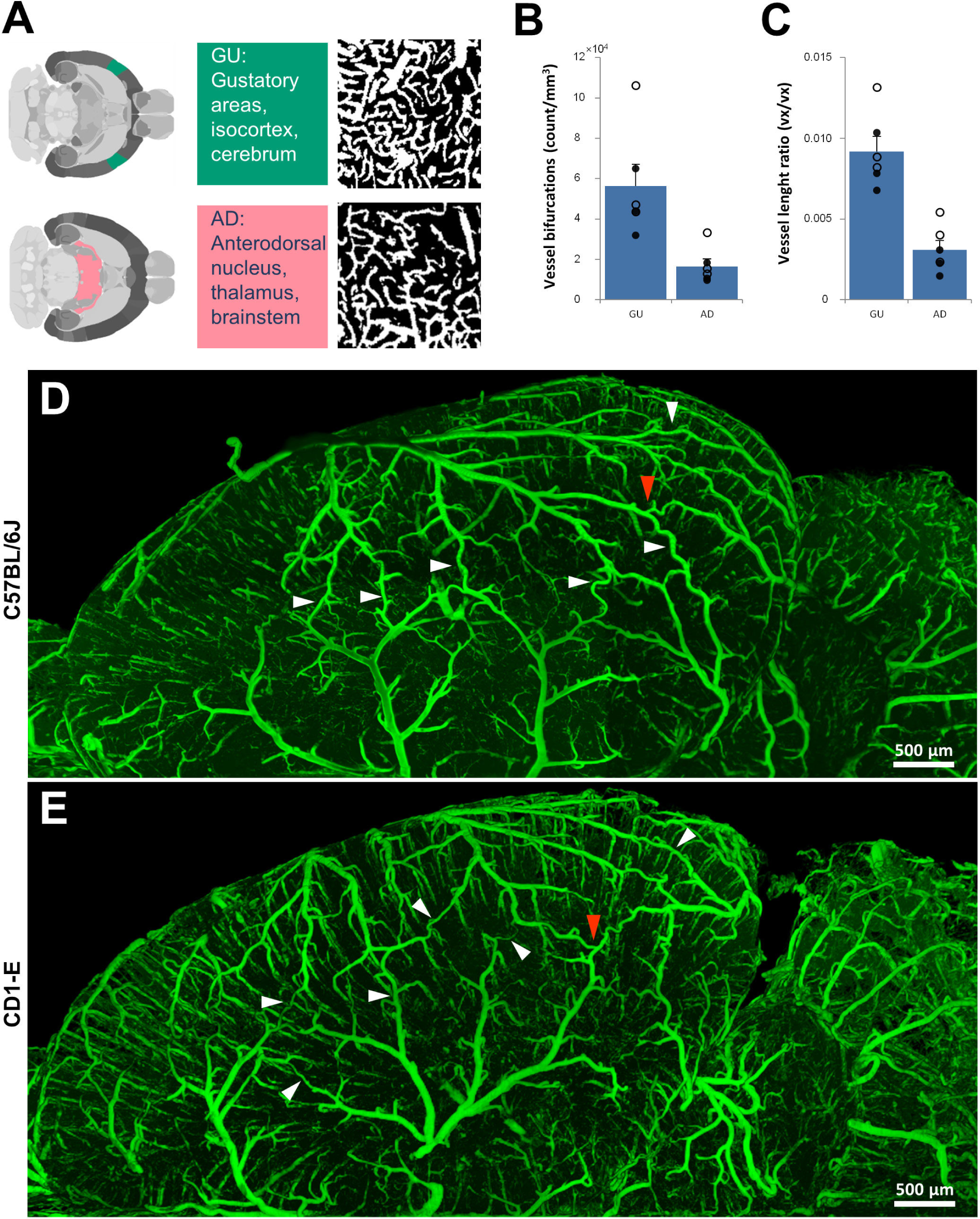
Exemplary quantitative analysis enabled by VesSAP. **A**, Maximum intensity projections representative patches from the Gustatory areas (GU) and Anterodorsal nucleus (AD) segmentation (600 x 600 x 33 µm). **B,C**, Quantification of the normalized bifurcation count and normalized vessel length ratio for the AD and GU clusters. In all graphs the black data points originate from the C57BL/6J samples and the white data points from the CD1-E. Mean values are given with error bars representing the standard error of the mean. **D,E**, Images of the vasculature in a representative C57BL/6J (**D**) and a CD1-E mouse (E) where the white arrowheads indicate the numerous anastomoses between the major arteries. Direct vascular connections between the medial cerebral artery (MCA), the anterior cerebral artery (ACA) and the posterior cerebral artery (PCER) are indicated by red arrowheads.

Importantly, our findings on vasculature length are in line with the predictions in the literature obtained in small volumes, confirming the robustness of our method. For example, previous studies quantifying small patches estimated the density of cortical blood vessels as 0.922 m/mm^3^, 0.444 m/mm^3^ or 0.471 m/mm^3^ in the cortex^12,17,37^. Using VesSAP, calculating centerline density as described above, and accounting for the 30 % isotropic tissue shrinkage in DISCO clearing^38^, we found an average vascular density of 0.473 ± 0.161 m/mm^3^ over the whole mouse cortex.

## DISCUSSION

Researchers have extensively worked towards examining the cerebral vasculature at the complete brain scale. This is particularly relevant, because current theories for many cardiovascular, neurodegenerative and metabolic disease pathologies include the capillaries, which can reflect the earliest symptoms. Thus, a method to robustly image, segment and analyze the complete cerebral vasculature of the mouse brain has been much needed. Previous studies were restricted: either they did not have sufficient resolutions to visualize all vessels including tiny capillaries such as MRI and microCT imaging modalities^39-41^, or once capillaries are imaged by high resolution fluorescent microscopy, the analysis could solely be done on small volumes^12^. Here we present VesSAP, a framework for unbiased statistical investigation of the complete vascular network in the intact adult mouse brain at the capillary level. We extracted the centerlines, bifurcation points and radius and assign them topographically to the Allen brain atlas to generate a reference map of the adult mouse brain vasculature. These maps can potentially be used to model synthetic cerebrovascular networks^42,43^. More advanced metrics to describe the vasculature and networks, for example global Strahler values, network connectivity and local statistics on bifurcation angles and vascular shape can be extracted using our method. Furthermore, the centerlines and bifurcation points can be interpreted as the edges and nodes for building a full vascular network graph, offering unprecedented means for studying local and global properties of the cerebrovascular network in the future.

Several methods have been proposed to label the cerebral vasculature of the mouse CNS based on one dye. Here, we employed two different dyes for complementary staining of the blood vessels, which are based on different mechanisms. The WGA binds to the glycocalyx of the endothelial lining of the blood vessels, but it can miss some segments of the large vessels. To circumvent this, we injected the EB dye into the mice 12 hours before WGA perfusion *in vivo*, allowing its long-term circulation to mark large vessels under physiological conditions. The combination of these dyes in this study enabled a wide dynamic contrast. This strategy has been proven quite beneficial for the segmentation of the complete cerebral vasculature of C57BL/6J and CD1-E strains as we showed here.

Our FCN segmentation architecture outperforms the current state-of-the-art methods significantly (**Fig. 3B**). Importantly, VesSAP is one to two orders of magnitude faster than the state-of-the-art automation which has a far lower accuracy, and more than 350 times faster than a human annotator. Our inter-annotator experiment validates that our model reaches human level performance (**Fig. 3C**). However, from a medical imaging and machine learning perspective the scores, especially dice, precision and recall are low compared to some other segmentation tasks in medical imaging^44-46^.

We attribute this mainly to the nature of a vascular network. Vessels are long but thin tubular shapes. In our images the radius of the capillaries (about 3 µm) is in the range of our voxel resolution, therefore, a segmentation of the correct thickness down to a single pixel is difficult. This inconsistency with the label does not pose a major problem for our main task to segment the whole vasculature and extract features; however it introduces a significant reduction to the F1Score, which is frequently used metric in machine learning tasks. The goal here is to segment a complete vasculature of the brain, to enable us to extract vascular features such as centerlines and bifurcation points. Therefore, introduction of a new metric in the future would be needed for this type of data. This metric should weight the correct detection of the vessels more and allow the outer wall of a blood vessel to be in a certain range of distance from the centerline of that vessel. The inter-annotator experiment and the comparison to our segmentation yielded insight in the quality of the segmentation. The human annotator segmentation accuracy and F1-Score were not superior compared to our model and the annotators disagreed substantially emphasizing the strength of our segmentation.

The proposed segmentation concept is based on a transfer learning approach, where we trained our data on a synthetic dataset and only refined it on a real labeled dataset 1.7 % of the size of the synthetic dataset. We consider this a major advantage compared to approaches that train from scratch. The model should generalize very well to other datasets, where other scientist would only need a small labeled dataset to achieve good segmentation performance. The pertinence of our approach should go well beyond vessel datasets, and could find applications for many other imaging tasks, for example tracing neurons.

Based on our vascular reference map new properties can be discovered and biological models can be confirmed faster. Here, we found that a substantial difference in vascularization and bifurcation density exists across the regions of the Allen brain atlas. Furthermore, we found that intracranial anastomoses between the anterior cerebral artery, middle cerebral artery and the posterior cerebral artery which are known from C57BL/6J^35,47^, are also present in albino CD1-Elite strain. This is opposed to the BALB/c albino mouse where the absence of collaterals has been described^35^. To our best knowledge this is the first evidence for high collateral density in albino CD1-Elite mice. This finding is important because existence of such collateral vessels between large vessels can significantly alter the outcome of the ischemic stroke lesion as the blood deprived brain regions from the occlusion of a large vessel could be compensated by the blood supplies coming from the collateral extensions from other large vessels^35,48,49^. Thus, our VesSAP method enables unbiased quantification of vascular anatomy in intact mouse brains and can lead to the discovery of previously overlooked anatomical knowledge.

In conclusion, VesSAP is the first scalable and automated machine learning-based method to analyze complex imaging data coming from the cleared intact mouse brains. It outperforms all previous methods of vessel segmentation and achieves a human level of accuracy several orders of magnitude faster. Thus, we foresee that our new method will accelerate the applications of tissue clearing in particular for the studies assessing the brain vasculature.

## METHODS

### Tissue preparation

The animals were housed under a 12/12 hr light/dark cycle. The animal experiments were conducted according to institutional guidelines (Klinikum der Universität München/Ludwig Maximilian University of Munich), after approval of the ethical review board of the government of Upper Bavaria (Regierung von Oberbayern, Munich, Germany), and in accordance with the European directive 2010/63/EU for animal research. For this study we injected 150 μl (2% V/V% in saline) of Evans blue dye (Sigma-Aldrich, E2129) intra-peritoneally into three C57BL/6J and CD1-Elite male, 3 months old mice (n=3 per group). After 12 hrs of postinjection time, we anaesthetized the animals with a triple combination of MMF (i.p.; 1 ml per 100 g body mass for mice) and opened their chest for transcardial perfusion. The following media was supplied by a peristaltic pump set to deliver 8 ml/min volume: 150 μl wheat germ agglutinin conjugated to Alexa 594 dye (ThermoFisher Scientific, W11262) and 15 ml PBS 1x and 15 ml 4% PFA.

After perfusion, the brains were extracted and incubated into 3DISCO clearing solutions as described by Ertürk et al.^21^. Briefly, we immersed them in a gradient of tetra-hydrofuran (Sigma-Aldrich, 186562): 50 vol%, 70 vol%, 80 vol%, 90 vol%, 100 vol% (in distilled water), and 100 vol% at 25 °C for 12 h each step. At this point we modified the protocol to incubate the samples in tert-Butanol incubation for 12 hrs at 35 °C followed by immersion in dichloromethane (Sigma-Aldrich, 270997) for 12 hrs at room temperature and finally incubation with the refractive index matching solution BABB (benzyl alcohol + benzyl benzoate 1:2; Sigma-Aldrich, 24122 and W213802), for at least 24 hrs at room temperature until transparency was achieved. Each incubation step was carried out on a laboratory shaker.

### Imaging of the cleared samples

We captured the optical section images with a 4× objective lens (Olympus XLFLUOR 340) equipped with an immersion corrected dipping cap mounted on a LaVision UltraII microscope. For 20× imaging, we used Zeiss CLARITY objective (Clr Plan-Neofluar, NA 1.0). The images were taken in 16 bit precision, which results in a resolution of 1.625 μm on the XY axes. The brain structures were visualized by the Alexa 594 and Evans blue fluorescent dyes at 561 and 640 nm excitation respectively. In z-dimension we took the sectional images in 3 μm steps from the right and left sides. To reduce defocus, which derives from the Gaussian shape of the beam we used a 12 step sequential shifting of the focal position of the light sheet per plane and side. The thinnest point of the light sheet was 5 μm.

### Reconstruction of the whole brain datasets from the tiling volumes

We stitched the acquired volumes using TeraStitcher’s automatic global optimization function (v1.10.3). We produced volumetric intensity images of the whole brain considering each channel separately. Next, we generated isotropic datasets because the registration and successive processing steps were more robust on isotropic datasets, therefore we downsampled the reconstructed 3D vascular datasets in XY dimensions to 3 × 3 × 3 μm resolution.

### Deep learning network architecture

Here we introduced a deep 3D fully convolutional network (FCN) for segmentation of our blood vessel dataset. The networks general architecture consists of 4 convolutional layers followed by a sigmoid activation layer, see **Fig. 3A**. Generally, the input layer is designed to take n images as an input. In our implemented case, the input to the first layer of the network are n=2 images of the same brain, which have been stained differently, see **Fig. 3A**. To specifically account for the general class imbalance (much more tissue background than vessels) in our dataset, and the high false positive rates associated with the class imbalance, the following class balancing loss function with stable weights from Tetteh et al. is implemented, see **Equation II.1**. Here, L_1_ is a numerically stable class balancing loss function and the term L_2_ penalizes the network for false predictions. Y_+_ and Y_-_ represent the foreground and background classes respectively, P(y_j_=k| X;W) is the probability that the voxel j in volume X belongs to class k given the volume X and network weights W. Y_f+_ and Y_f-_ represent the false positive and false positive predictions respectively at each training iteration.

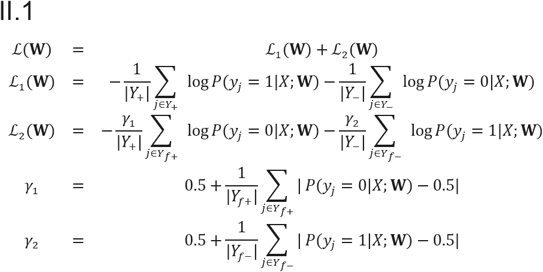

The 3D convolutional operations in this network are implemented as sparse crosshair filters to reduce memory consumption and speed up the computation, for a graphical representation see **Supporting Fig. 4**. Tetteh et al. showed that by using this operation a faster computation is achieved without undermining the prediction accuracy^25^. The crosshair filter works by separating a full 3D kernel into 3 orthogonal 2D kernels. Those kernels are applied to the volume at every layer of the network.

The networks training is driven by a stochastic gradient descent function without a regularization. A prediction or segmentation with a trained model takes a volumetric image of arbitrary size and outputs an estimated probabilistic segmentation of the input images size. The algorithms have been implemented using the THEANO framework^50^. They are trained and tested on a NVIDIA Quadro P5000 GPU and on machines with 64GB and 500GB RAM respectively.

### Transfer learning

Typically, medical imaging tasks are aggravated by scarce and very scarce data availability. The proposed transfer learning approach, aims to account for the scarce labeled data by pre-training our models on a synthetic dataset and refining them on a small training set of interest^51^. Our approach pre-trains a two channel version of DeepVesselNet on a synthetically generated dataset^52,53^, with the goal to learn specific vascular shaped image patterns. The pre-training is carried out on a dataset of 136 volumes of a size of 325 × 304 × 600 pixels. While pre-training we applied a learning rate of 0.01 and a decay of 0.99 which we applied after every 200 iterations. Finally the pre-trained model is fine-tuned by re-training on a real microscopic dataset consisting of eleven volumes with a size of 500 × 500 × 50 pixels, which were manually annotated by the expert who imaged the data and further verified by two additional experts. All volumes are processed in smaller sub-patches by our network. This enables us to process volumes of arbitrary sizes and dimensions. The data we use in the fine-tuning step accumulates to 1.7% of the synthetic datasets voxel volume and solely 0.02% of the voxel volume of a single brain image. For the fine-tuning step we utilized a learning rate of 0.0001 and a decay of 0.98, which we applied after every 10 iterations.

Our training set consist of eleven volumetric images from two mice brains, the test and validation set consists of four patches from two different brains. Each patch consists of a volume of 500 × 500 × 50 pixels. We chose independent brains to guarantee generalizability. The patches are processed and predicted in 25 small sub-patches. We cross-test on our test and validation set by rotating these four-fold. In every rotation our validation set consists of 3 patches and our test set of one patch. To prevent an overfitting of our model we chose the validation and test set from two brains. One from the CD1-E and one from the C57BL/6J strain. We choose the lowest log loss on our validation set to be our model selection point (see **Supporting Fig. 4a**). We report an average F1-Score of 0.76 ± 0.01, an average accuracy of 0.93 ± 0.01, an average precision of 0.79 ± 0.02 and an average recall of 0.73 ± 0.02 on our test sets. All scores are given with a 1σ standard deviation. On average our model reached the model selection point after 45 epochs of training.

### Pre-processing of segmentation

The pre-processing represents a significant factor for the overall success of the training and segmentation. The intensity distribution among the brains and among brain regions differs substantially. To account for the intensity distributions, two preprocessing strategies have been applied successively.

a. High-cut filter: In this step the intensities x above a certain threshold, c which is defined by an individual percentile for each volume is set to that threshold. Next, they were normalized by f(x).

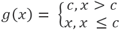
b. Normalization of intensities: The original intensities were normalized to the range of 0 to 1, where x is the pixel intensity and X are all intensities of the volume.

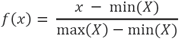

### Inter-annotator experiment

To compare VesSAP’s segmentation to a human level annotation we implemented an inter-annotator experiment. For this experiment we determined a gold standard label for two patches of 500 × 500 × 500 pixels from a commission of three experts, including the expert who imaged our data and is therefore most familiar with the images. Next, we gave the two patches to 4 other experts to label the complete vasculature. The experts spend multiple hours to label each patch within the ImageJ and ITK-snap environment and were allowed to use their favored approaches to generate their best label. Finally, we calculated the accuracy and dice scores for the different raters, compared to the gold standard and compared them to the scores of our model.

### Feature extraction

In order to quantify the anatomy of the mouse brain vasculature we extracted descriptive features based on our segmentation. Later we registered them to the Allen brain atlas.

As features we extracted the centerlines, the bifurcation points and the radius of the segmented blood vessels. We consider those features to be independent from the elongation of the light sheet scans and the connectedness of the vessels due to staining, imaging and/or segmentation artefacts. We found the extracted features as a baseline.

Before extracting the centerlines we applied two cycles of binary erosion and dilation to remove false negative pixels within the volume of segmented vessels as those would induce false centerlines. Our centerline extraction is based on a 3D thinning algorithm as introduced by Lee et al.^54^. Based on the centerlines we extracted bifurcation points. A bifurcation is the branching point on a centerline where a larger vessel splits into two or more small vessels (see **Fig. 4A)**. In a network analysis context they are significant as they represent the nodes of a vascular network^55^. Furthermore, bifurcation points have significance in a biological context. In neurodegenerative diseases, capillaries are known to degenerate^56^, thereby significantly reducing the number of bifurcation points in an affected brain region compared to a healthy brain. To detect the bifurcation points an algorithm was implemented. The algorithm takes the centerlines as an input and calculates for every point on that centerline the surrounding centerline pixels to determine if a point is a centerline. The radius of a blood vessel is a key feature to describe vascular networks. The radius yields information about the flow and hierarchy of the vessel network^55^. The proposed algorithm calculates the Euclidean distance transform for every segmented pixel v to the closest background pixel b_closest_ (**Equation II.2**). Next, the distance transform matrix is multiplied with the 3D centerline mask equaling the minimum radius of the vessel around the centerline.

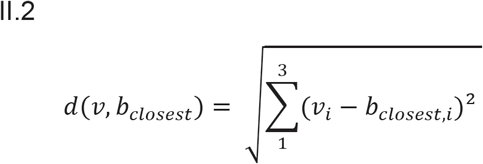

### Registration to the reference atlas

We used the average template, the annotation file and the latest ontology file (Ontology ID: 1) of the current Allen brain mouse atlas CCFv3 201710. Then we scaled the template and the annotation file up from 10 to 3 µm^3^ to match our reconstructed brain scans. After this we multiplied the left side of the (still symmetrical) annotation file with −1 so that the labels can be later assigned to the corresponding hemispheres. Next, the average template and the 3D vascular datasets were downsampled to 10% of their original size in each dimension to achieve a reasonably fast alignment. In the sake of the integrity of the extracted features, we aligned the template to each of the brain scans individually using a two-step rigid and deformable (B-Spline) registration and applied the transformation parameters to the full resolution annotation volume in 3 × 3 × 3 μm resolution. Subsequently we created masks for the anatomical clusters based on the current Allen brain atlas ontology.

### Statistics

Data collection and analysis were not performed blind to the strains. Data distribution was assumed to be normal, but this was not formally tested. All data values are given as mean ± SEM. Data were analyzed with standardized effect size indices (Cohen’s *d*)^57^ to investigate differences of vessel density, number of bifurcation points and radii between brain areas across the two mouse strains (n=3 per strain) and comparisons across brain areas in the pooled (n=6) dataset. Statistical analysis was performed using MATLAB.

### Data visualization

All volumetric datasets were rendered using Imaris, Arivis and ITK Snap.

## CODE AND DATA AVAILABILITY

VesSAP codes and data that we produced are publicly hosted online for easy adoption, including the imaging protocol, the data (original scans, registered atlas data), the trained algorithms, training data and a reference set of features describing the vascular network in all brain regions at the following address. Implementation of external libraries are available on request. http://DISCOtechnologies.org/VesSAP

## ACKNOWLEDGMENTS

This work was supported by the Vascular Dementia Research Foundation, Synergy Excellence Cluster Munich (SyNergy), ERA-Net Neuron (01EW1501A to A.E.), Fritz Thyssen Stiftung (A.E., Ref. 10.17.1.019MN), DFG (A.E., Ref. ER 810/2-1), NIH (A.E.), Helmholtz ICEMED Alliance (A.E.), and the German Federal Ministry of Edu-cation and Research via the Software Campus initiative (O.S.). Furthermore, NVIDIA supported this work with a Titan XP via the GPU Grant Program. M.I.T is member of Graduate School of Systemic Neurosciences (GSN), Ludwig Maximilian University of Munich.

## AUTHOR CONTRIBUTIONS

M.I.T. performed the tissue processing, clearing and imaging experiments. M.I.T and K.V. developed the whole brain staining protocol. M.I.T. stitched and assembled the whole brain scans. V.E. generated the synthetic vascular training dataset. J.C.P, G.T. and O.S developed the deep learning architecture, trained the models and performed the quantitative analyses. M.I.T. annotated the data. M.D. and M.D. helped with data interpretation. B.M, M.P. and G.T. provided guidance in developing the deep learning architecture and helped with data interpretation. A.E., M.I.T. and J.C.P. wrote the manuscript. All the authors edited the manuscript. A.E. initiated and led all aspects of the project.

## CONFLICT OF INTEREST STATEMENT

The authors declare that the research was conducted in the absence of any commercial or financial relationships that could be construed as a potential conflict of interest.

## VIDEO LEGENDS

### Supporting Video 1

Visualization of a representative CD1-E mouse brain by VesSAP showing the data quality.

### Supporting Video 2 (VR optimized viewing)

The whole mouse brain shown in Video 1 has been rendered for virtual reality (VR) viewing using Arivis InViewR. The immersed VR view shows the quality of VesSAP segmentation. We propose that scientific VR videos coming from large cleared samples could be a helpful tool for scientists to explore the data in a 3D interactive way. VR videos might also be used for educational purposes as they can be viewed on smart phones and other available VR devices. Please check the link for more information regarding how to view this VR video: **http://DISCOtechnologies.org/VesSAP/#VR**

### Supporting Video 3

Segmentation and features demonstration on a subset of the whole dataset. VesSAP enables reliable segmentation (red) and feature extraction (bifurcation points and centerlines, green and cyan) down to the capillary-level from the imaging data (grey).

### Supporting Video 4

Whole brain data registered to the Allen adult brain atlas. The video shows the alignment accuracy and segmentation overlaid.

### Supporting Video 5

Substack of the whole brain data registered to the Allen adult brain atlas. This video reveals the full resolution segmentation on a small set of the brain scans data.

---

**Supporting figure 1:**
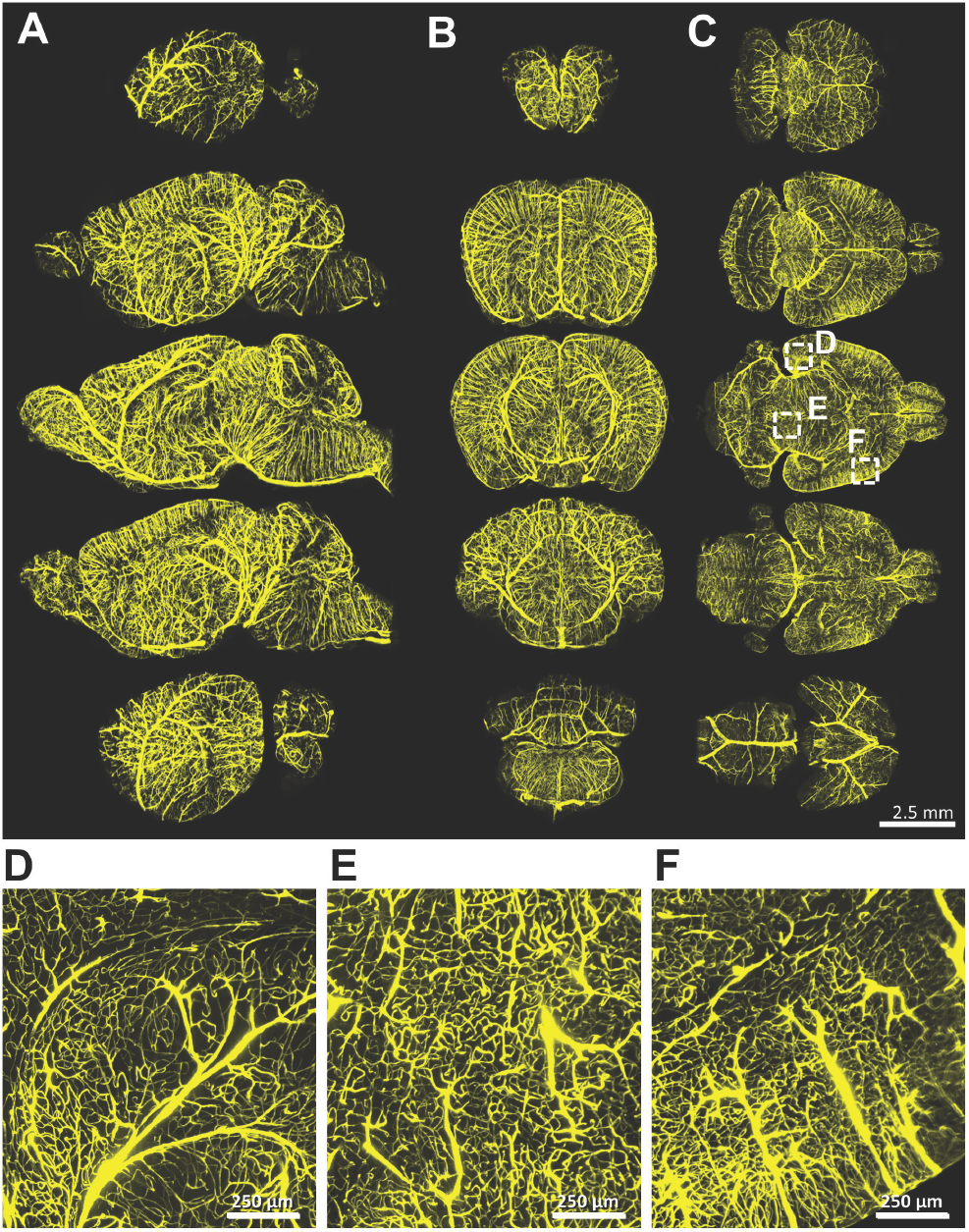
Vasculature is stained homogenously throughout all brain regions. **A**, Sagittal maximum intensity projections. **B**, Coronal maximum intensity projections. **C**, Axial maximum projections. **D-F**, Zoom-ins where capillary level staining is evident.

**Supporting figure 2:**
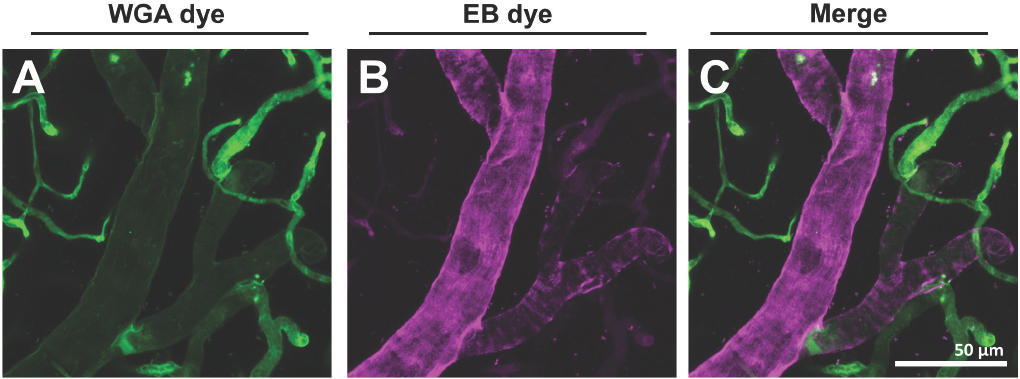
Confocal microscopy confirms that the neurovasculature is stained in a complimentary way. **A,B**, Maximum intensity projection of the WGA and the EB signal respectively. **C**, Merge of the two signals shows that capillaries are predominantly stained with WGA whereas EB shows strong staining of major blood vessels.

**Supporting figure 3:**
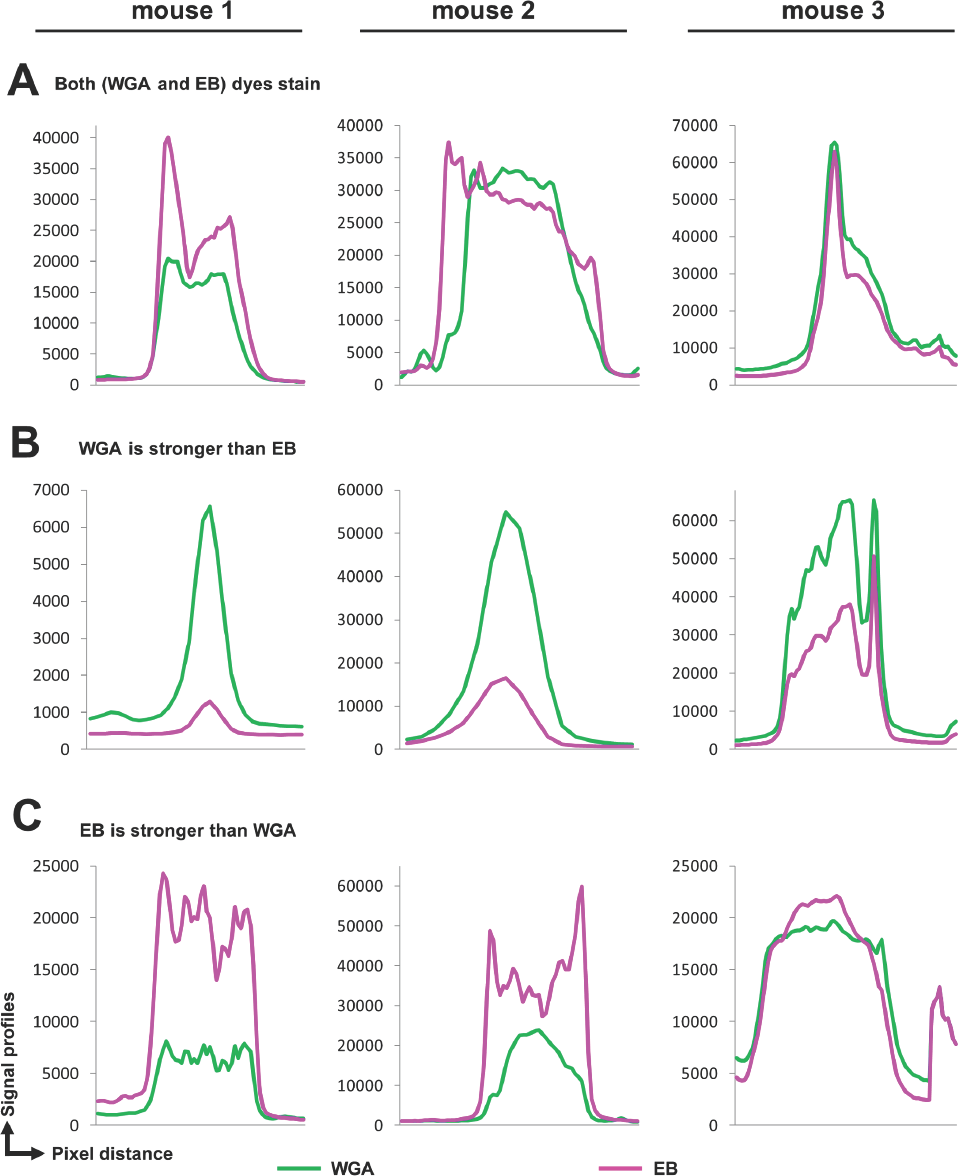
Raw signal intensity distribution along line profiles across stained vessels for three animals. Both dyes stain the vasculature with a complimentary SNR. For some vessels the SNR of both channels are similar (**A**), whereas for other vessels the EB or WGA channels have a substantially higher SNR compared to the other (**B**) and (**C**). These graphs quantitatively describe the SNR enhancements owing to double dye staining strategy.

**Supporting figure 4:**
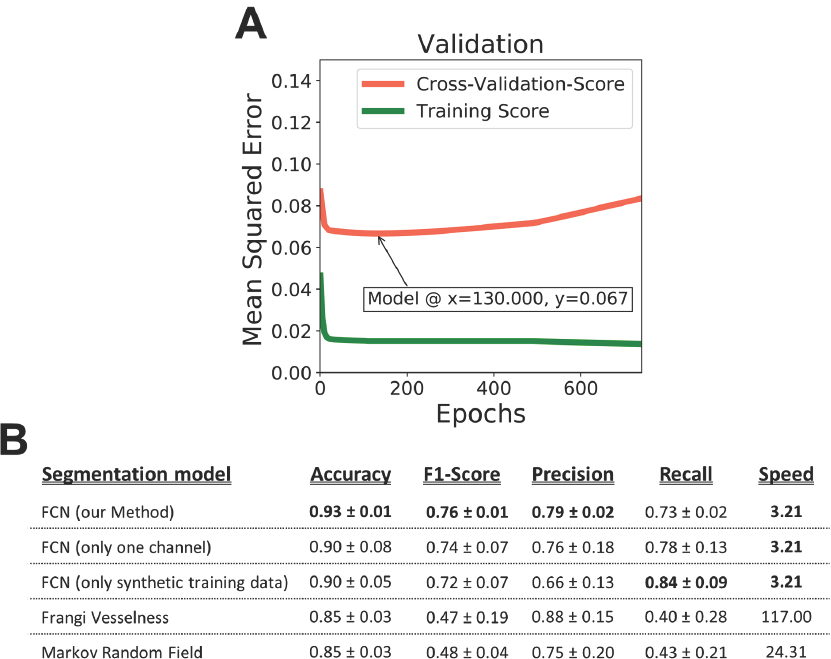
Details of VesSAP performance. **A**, Averaged validation performance and model selection point on the mean squared error metric. **B**, Evaluation metrics: accuracy, F1-Score, precision, recall and speed (for one image volume of 500 × 500 × 50 pixels) of the different models for segmentation.

**Supporting figure 5:**
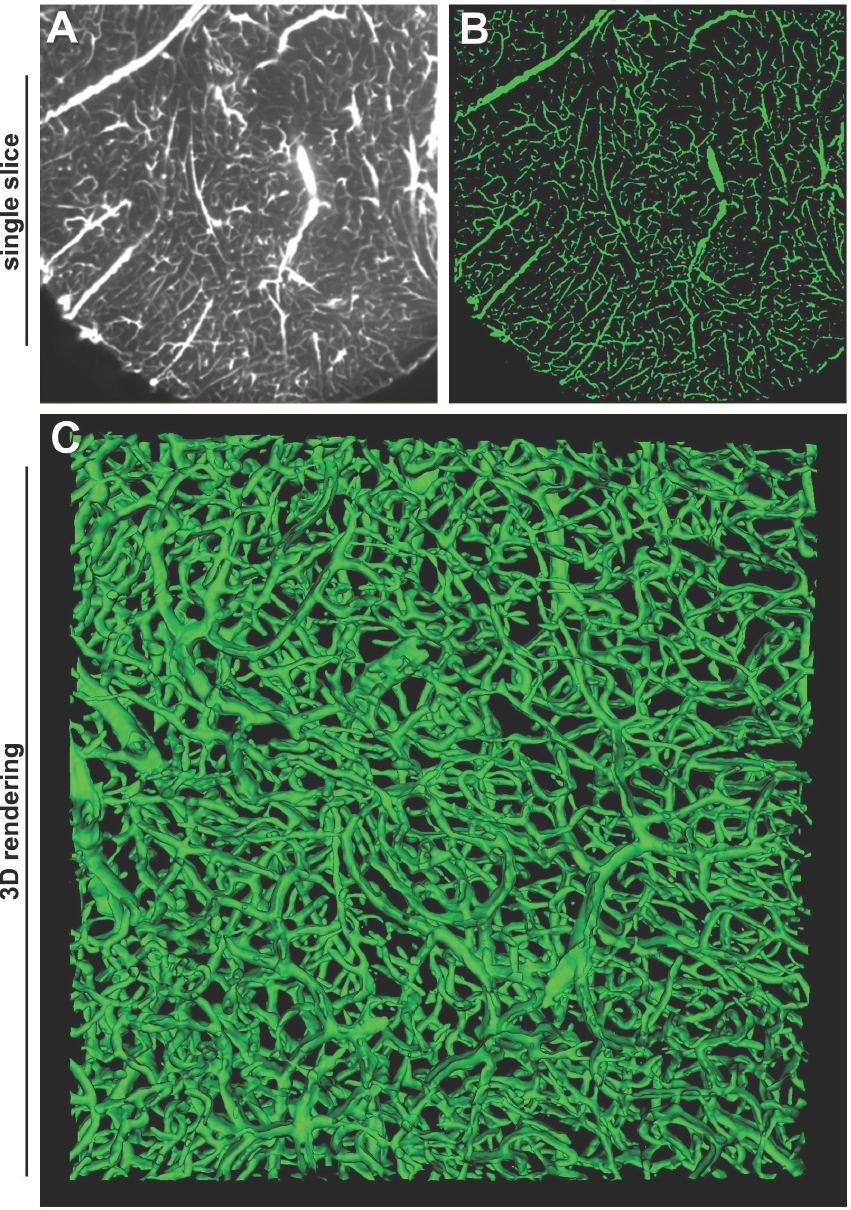
Details of the segmentation quality by VesSAP. **A,B**, Side by side slices of the raw lectin channel image and the segmentation (green). **C**, 3D rendering of a small brain patch showcasing connected capillaries.

**Supporting figure 6:**
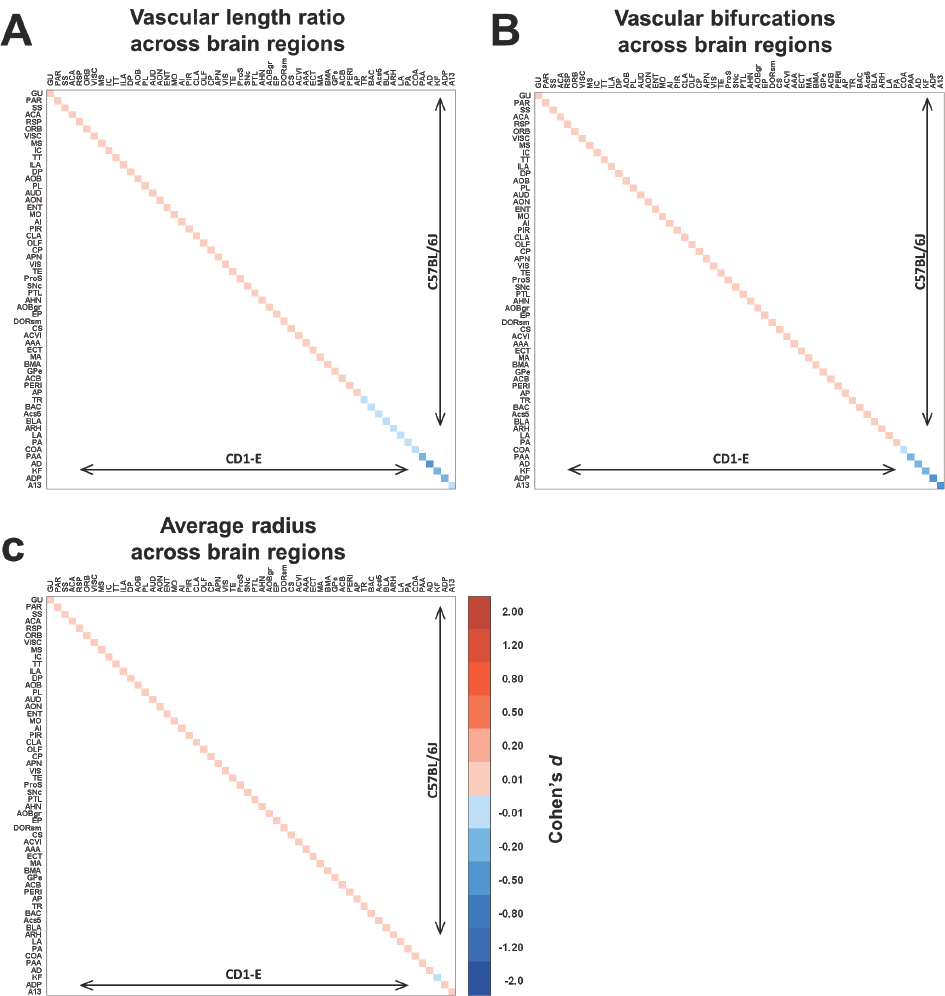
Inter-strain comparison of the features of the vascular network in the C57BL/6J and CD1-E mice using Cohen’s *d* method. **A-B**, Normalized vessel and bifurcations density matrices show small differences on the level of strains respectively. **C**, Distribution of average radius across brain regions in the two strains. It shows a mainly homogenous pattern, most probably governed by the high amount of capillaries in the vascular network. For the full list of abbreviations refer to the Supporting Table 1. The extracted numerical features are in Supporting Tables 2-4.

**Supporting table 1:**
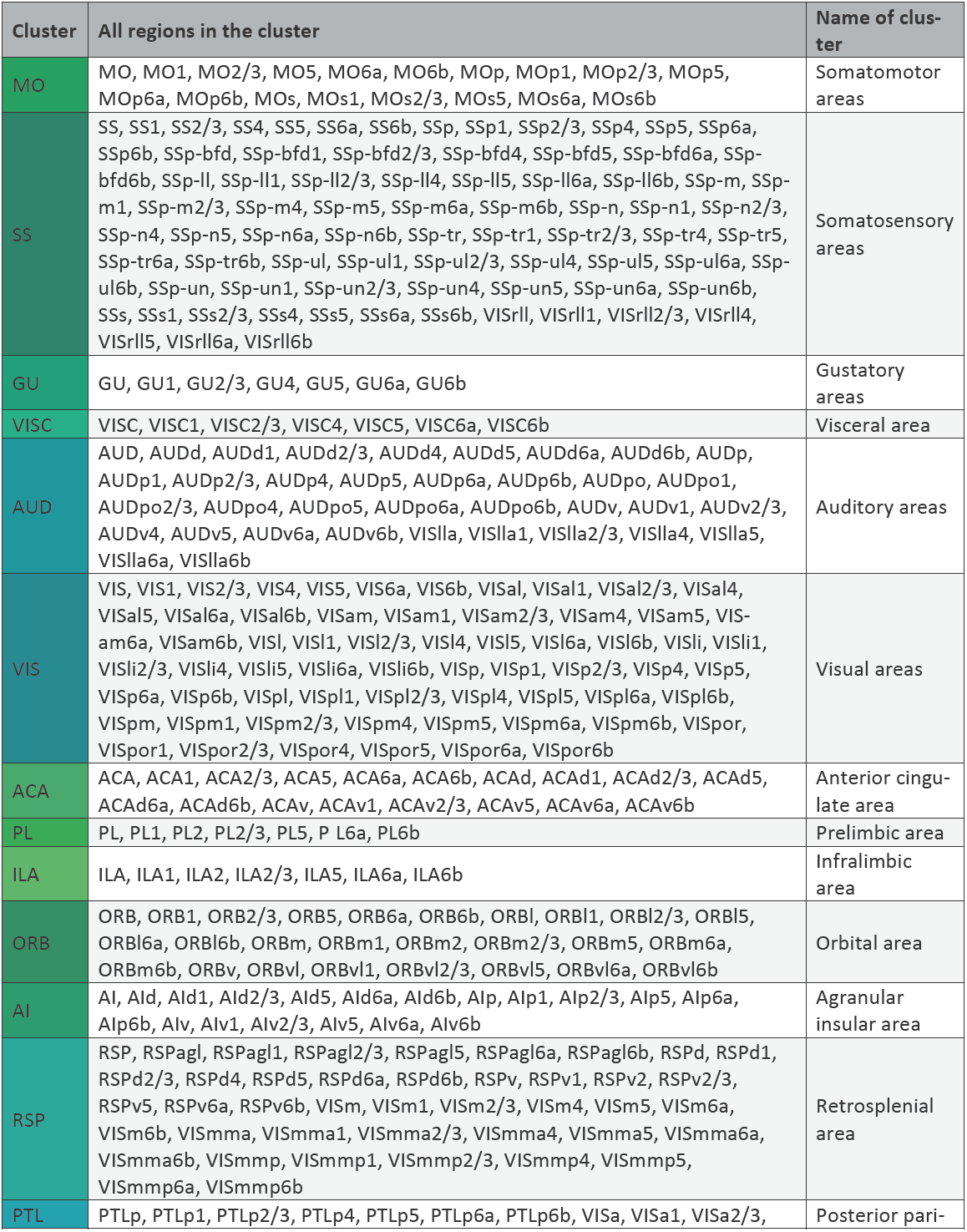
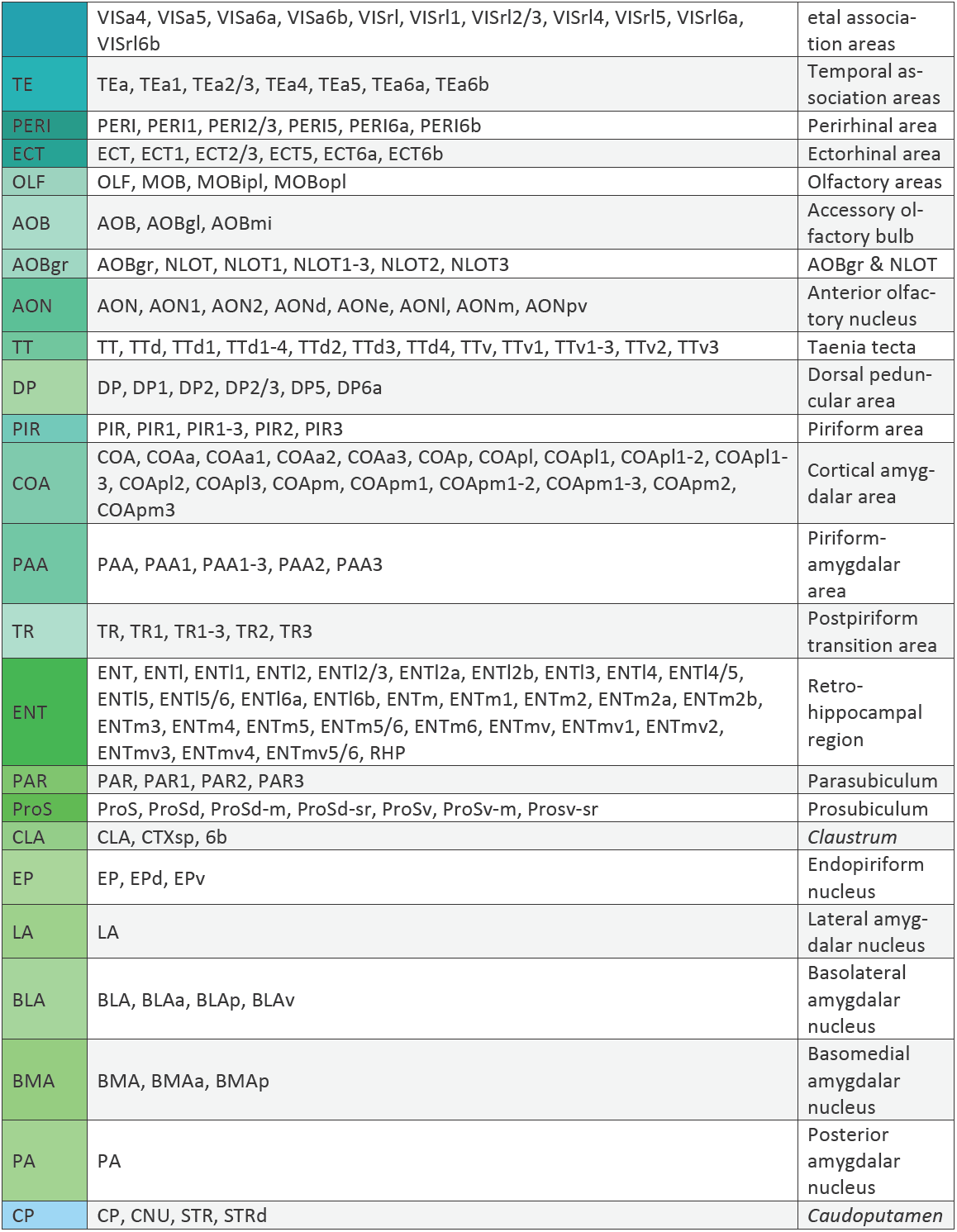
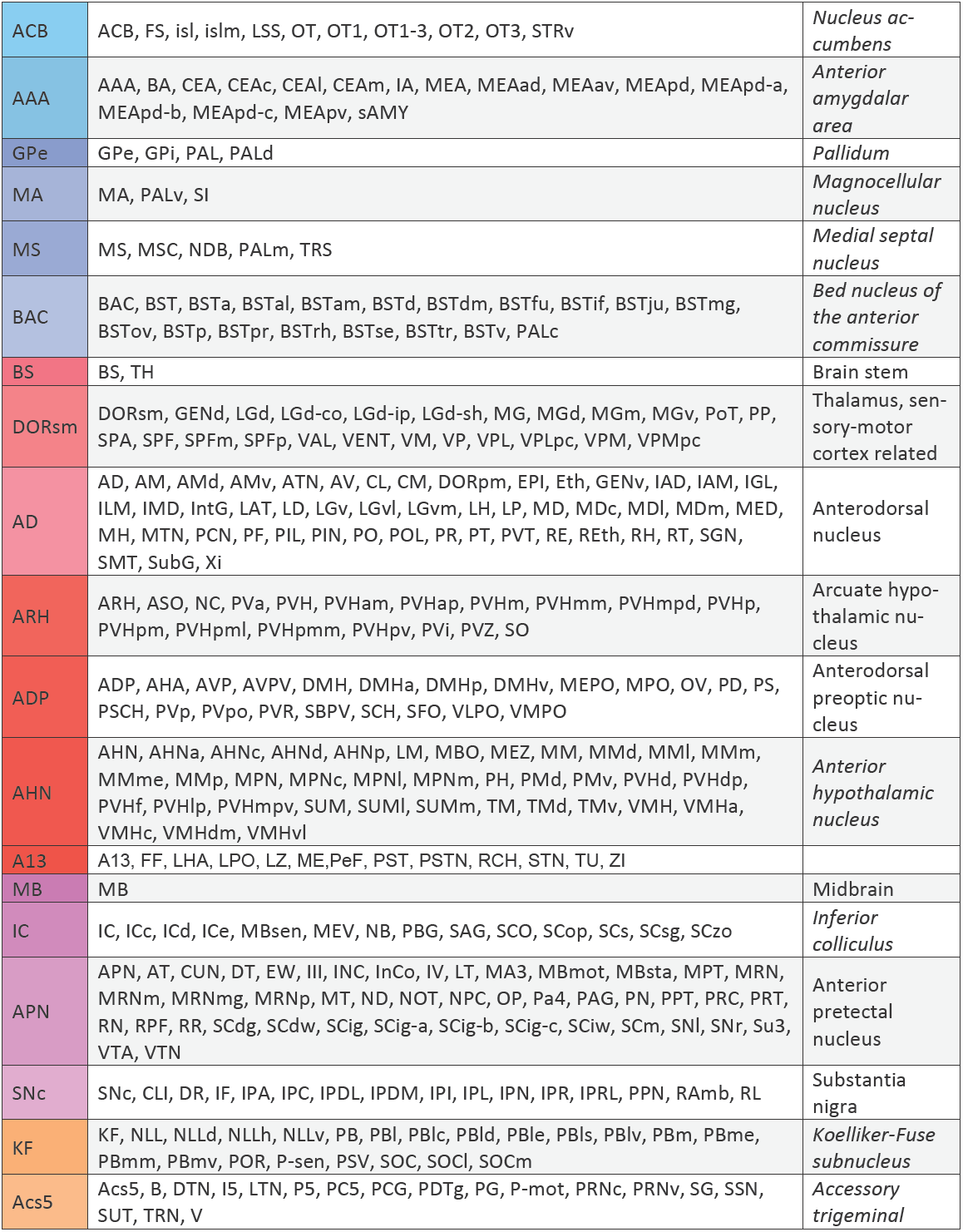
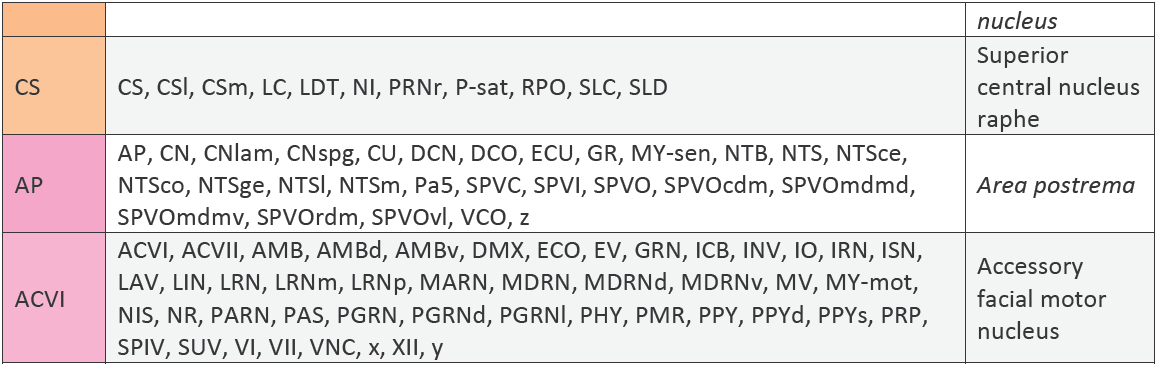
List of anatomical clusters and all the brain regions that they represent according to the current Allen adult mouse brain atlas ontology.

**Supporting table 2:** Quantification of the vascular density in the cleared C57BL/6J and CD1-E samples. Units are *voxel / voxel*.

| Cluster | BL6#2 | BL6#4 | BL6#5 | CD1#15 | CD1#41 | CD1#42 |
| --- | --- | --- | --- | --- | --- | --- |
| MO | 0.0093572 | 0.0059082 | 0.0047905 | 0.0052599 | 0.0074767 | 0.0056309 |
| SS | 0.0115752 | 0.0093265 | 0.0081327 | 0.0070534 | 0.0112136 | 0.0066858 |
| GU | 0.0103486 | 0.0078052 | 0.0067535 | 0.0088425 | 0.0131467 | 0.0082024 |
| VISC | 0.0094285 | 0.0077694 | 0.0060486 | 0.0062264 | 0.0122675 | 0.0053494 |
| AUD | 0.0076337 | 0.0058136 | 0.0059237 | 0.0069463 | 0.0075269 | 0.0061430 |
| VIS | 0.0076265 | 0.0050208 | 0.0054900 | 0.0043612 | 0.0069535 | 0.0042240 |
| ACA | 0.0108309 | 0.0070736 | 0.0060967 | 0.0081567 | 0.0111447 | 0.0089168 |
| PL | 0.0102455 | 0.0064844 | 0.0046423 | 0.0051858 | 0.0085280 | 0.0058823 |
| ILA | 0.0108894 | 0.0047988 | 0.0028369 | 0.0077173 | 0.0081559 | 0.0086919 |
| ORB | 0.0137248 | 0.0064118 | 0.0058307 | 0.0060286 | 0.0092827 | 0.0067783 |
| AI | 0.0093757 | 0.0058920 | 0.0049804 | 0.0049596 | 0.0078331 | 0.0047837 |
| RSP | 0.0115985 | 0.0091914 | 0.0062429 | 0.0053676 | 0.0109665 | 0.0055400 |
| PTL | 0.0045193 | 0.0048831 | 0.0052986 | 0.0048463 | 0.0068559 | 0.0044541 |
| TE | 0.0053130 | 0.0047694 | 0.0046928 | 0.0055785 | 0.0070274 | 0.0048437 |
| PERI | 0.0051592 | 0.0033750 | 0.0046063 | 0.0032898 | 0.0055644 | 0.0027928 |
| ECT | 0.0048977 | 0.0037841 | 0.0043316 | 0.0040392 | 0.0063336 | 0.0034538 |
| OLF | 0.0091333 | 0.0028067 | 0.0056325 | 0.0038479 | 0.0104136 | 0.0027701 |
| AOB | 0.0102001 | 0.0074197 | 0.0073297 | 0.0036032 | 0.0089601 | 0.0040691 |
| AOBgr | 0.0066158 | 0.0071247 | 0.0058011 | 0.0026772 | 0.0056548 | 0.0024047 |
| AON | 0.0119582 | 0.0032694 | 0.0047818 | 0.0050356 | 0.0085656 | 0.0055792 |
| TT | 0.0128843 | 0.0057909 | 0.0071700 | 0.0058206 | 0.0082569 | 0.0064400 |
| DP | 0.0107445 | 0.0037908 | 0.0039061 | 0.0071797 | 0.0082577 | 0.0080111 |
| PIR | 0.0093940 | 0.0062563 | 0.0065447 | 0.0042132 | 0.0074207 | 0.0037186 |
| COA | 0.0044112 | 0.0028131 | 0.0044865 | 0.0021680 | 0.0045812 | 0.0017700 |
| PAA | 0.0042952 | 0.0022902 | 0.0043477 | 0.0017543 | 0.0050305 | 0.0014458 |
| TR | 0.0058219 | 0.0029637 | 0.0044652 | 0.0027056 | 0.0056278 | 0.0022267 |
| ENT | 0.0063865 | 0.0051335 | 0.0046995 | 0.0042859 | 0.0144999 | 0.0036851 |
| PAR | 0.0114697 | 0.0091646 | 0.0076682 | 0.0078766 | 0.0113733 | 0.0070761 |
| ProS | 0.0076081 | 0.0038296 | 0.0033865 | 0.0050129 | 0.0076083 | 0.0047397 |

**Supporting table 2:** Quantification of the vascular density in the cleared C57BL/6J and CD1-E samples.
|  |  |  |  |  |  |  |
| --- | --- | --- | --- | --- | --- | --- |
| CLA | 0.0096760 | 0.0053913 | 0.0053636 | 0.0045884 | 0.0071508 | 0.0042252 |
| EP | 0.0078594 | 0.0046981 | 0.0047499 | 0.0035476 | 0.0062039 | 0.0031096 |
| LA | 0.0048305 | 0.0019873 | 0.0032737 | 0.0032051 | 0.0061426 | 0.0026785 |
| BLA | 0.0054087 | 0.0022658 | 0.0039579 | 0.0026897 | 0.0058530 | 0.0022209 |
| BMA | 0.0064536 | 0.0030861 | 0.0045863 | 0.0031615 | 0.0057182 | 0.0026441 |
| PA | 0.0042582 | 0.0035128 | 0.0024244 | 0.0031166 | 0.0051596 | 0.0025716 |
| CP | 0.0083593 | 0.0037503 | 0.0048630 | 0.0050527 | 0.0074828 | 0.0048447 |
| ACB | 0.0066948 | 0.0019018 | 0.0043521 | 0.0033112 | 0.0055023 | 0.0034835 |
| AAA | 0.0074127 | 0.0041995 | 0.0033715 | 0.0036295 | 0.0056303 | 0.0030950 |
| GPe | 0.0069966 | 0.0024761 | 0.0040185 | 0.0030260 | 0.0063187 | 0.0027351 |
| MA | 0.0085149 | 0.0021984 | 0.0047591 | 0.0025952 | 0.0062517 | 0.0024843 |
| MS | 0.0113546 | 0.0072848 | 0.0077730 | 0.0056466 | 0.0092087 | 0.0057410 |
| BAC | 0.0064581 | 0.0012580 | 0.0036531 | 0.0030268 | 0.0061399 | 0.0029772 |
| BS | 0.0048691 | 0.0024906 | 0.0035570 | 0.0051794 | 0.0088115 | 0.0050795 |
| DORsm | 0.0030760 | 0.0014435 | 0.0022406 | 0.0023438 | 0.0054218 | 0.0040204 |
| AD | 0.0056753 | 0.0022222 | 0.0033178 | 0.0027231 | 0.0057565 | 0.0026248 |
| ARH | 0.0035346 | 0.0013719 | 0.0019229 | 0.0018142 | 0.0033885 | 0.0042785 |
| ADP | 0.0080422 | 0.0043285 | 0.0048229 | 0.0034813 | 0.0069629 | 0.0031331 |
| AHN | 0.0032352 | 0.0012860 | 0.0019369 | 0.0007876 | 0.0016155 | 0.0044040 |
| A13 | 0.0057085 | 0.0087157 | 0.0037526 | 0.0076900 | 0.0131571 | 0.0079402 |
| MB | 0.0073036 | 0.0034578 | 0.0045716 | 0.0052940 | 0.0075160 | 0.0060563 |
| IC | 0.0069680 | 0.0038629 | 0.0047105 | 0.0045880 | 0.0078609 | 0.0041552 |
| APN | 0.0029680 | 0.0032348 | 0.0030604 | 0.0022715 | 0.0047944 | 0.0019137 |
| SNc | 0.0060860 | 0.0019211 | 0.0044188 | 0.0028324 | 0.0057872 | 0.0023751 |
| KF | 0.0076008 | 0.0018858 | 0.0044324 | 0.0033314 | 0.0085823 | 0.0028944 |
| Acs5 | 0.0066081 | 0.0025452 | 0.0046771 | 0.0036584 | 0.0039047 | 0.0032545 |
| CS | 0.0087011 | 0.0026383 | 0.0057563 | 0.0035171 | 0.0045107 | 0.0031765 |
| AP | 0.0066081 | 0.0025452 | 0.0046771 | 0.0036584 | 0.0039047 | 0.0032545 |
| ACVI | 0.0087011 | 0.0026383 | 0.0057563 | 0.0035171 | 0.0045107 | 0.0031765 |

**Supporting table 3:**
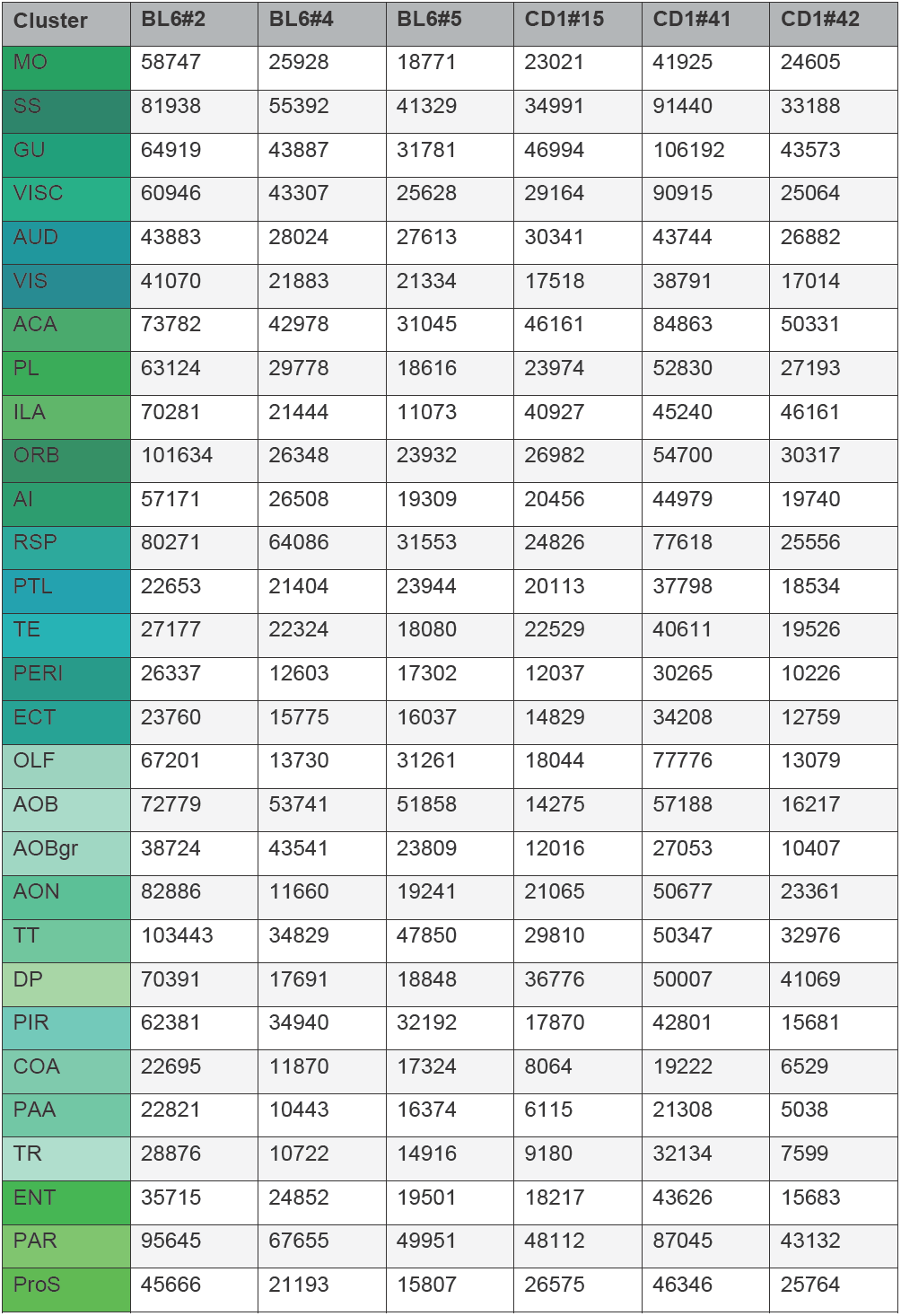
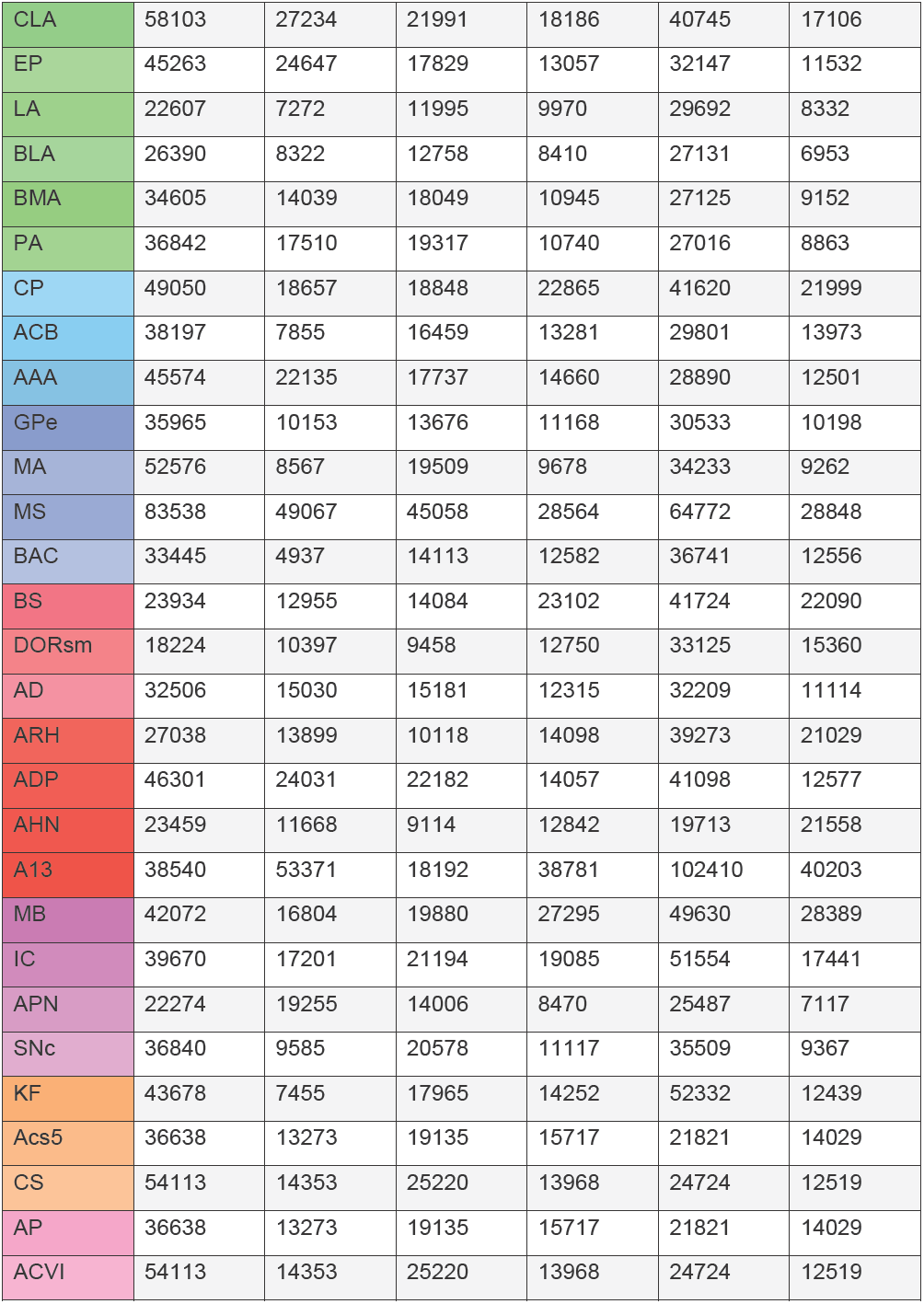
Quantification of the number of bifurcation points in the cleared C57BL/6J and CD1-E samples, units are *counts / mm*^*3*^.

| Cluster | BL6#2 | BL6#4 | BL6#5 | CD1#15 | CD1#41 | CD1#42 |
| --- | --- | --- | --- | --- | --- | --- |
| MO | 58747 | 25928 | 18771 | 23021 | 41925 | 24605 |
| SS | 81938 | 55392 | 41329 | 34991 | 91440 | 33188 |
| GU | 64919 | 43887 | 31781 | 46994 | 106192 | 43573 |
| VISC | 60946 | 43307 | 25628 | 29164 | 90915 | 25064 |
| AUD | 43883 | 28024 | 27613 | 30341 | 43744 | 26882 |
| VIS | 41070 | 21883 | 21334 | 17518 | 38791 | 17014 |
| ACA | 73782 | 42978 | 31045 | 46161 | 84863 | 50331 |
| PL | 63124 | 29778 | 18616 | 23974 | 52830 | 27193 |
| ILA | 70281 | 21444 | 11073 | 40927 | 45240 | 46161 |
| ORB | 101634 | 26348 | 23932 | 26982 | 54700 | 30317 |
| AI | 57171 | 26508 | 19309 | 20456 | 44979 | 19740 |
| RSP | 80271 | 64086 | 31553 | 24826 | 77618 | 25556 |
| PTL | 22653 | 21404 | 23944 | 20113 | 37798 | 18534 |
| TE | 27177 | 22324 | 18080 | 22529 | 40611 | 19526 |
| PERI | 26337 | 12603 | 17302 | 12037 | 30265 | 10226 |
| ECT | 23760 | 15775 | 16037 | 14829 | 34208 | 12759 |
| OLF | 67201 | 13730 | 31261 | 18044 | 77776 | 13079 |
| AOB | 72779 | 53741 | 51858 | 14275 | 57188 | 16217 |
| AOBgr | 38724 | 43541 | 23809 | 12016 | 27053 | 10407 |
| AON | 82886 | 11660 | 19241 | 21065 | 50677 | 23361 |
| TT | 103443 | 34829 | 47850 | 29810 | 50347 | 32976 |
| DP | 70391 | 17691 | 18848 | 36776 | 50007 | 41069 |
| PIR | 62381 | 34940 | 32192 | 17870 | 42801 | 15681 |
| COA | 22695 | 11870 | 17324 | 8064 | 19222 | 6529 |
| PAA | 22821 | 10443 | 16374 | 6115 | 21308 | 5038 |
| TR | 28876 | 10722 | 14916 | 9180 | 32134 | 7599 |
| ENT | 35715 | 24852 | 19501 | 18217 | 43626 | 15683 |
| PAR | 95645 | 67655 | 49951 | 48112 | 87045 | 43132 |
| ProS | 45666 | 21193 | 15807 | 26575 | 46346 | 25764 |

|  |  |  |  |  |  |  |
| --- | --- | --- | --- | --- | --- | --- |
| CLA | 58103 | 27234 | 21991 | 18186 | 40745 | 17106 |
| EP | 45263 | 24647 | 17829 | 13057 | 32147 | 11532 |
| LA | 22607 | 7272 | 11995 | 9970 | 29692 | 8332 |
| BLA | 26390 | 8322 | 12758 | 8410 | 27131 | 6953 |
| BMA | 34605 | 14039 | 18049 | 10945 | 27125 | 9152 |
| PA | 36842 | 17510 | 19317 | 10740 | 27016 | 8863 |
| CP | 49050 | 18657 | 18848 | 22865 | 41620 | 21999 |
| ACB | 38197 | 7855 | 16459 | 13281 | 29801 | 13973 |
| AAA | 45574 | 22135 | 17737 | 14660 | 28890 | 12501 |
| GPe | 35965 | 10153 | 13676 | 11168 | 30533 | 10198 |
| MA | 52576 | 8567 | 19509 | 9678 | 34233 | 9262 |
| MS | 83538 | 49067 | 45058 | 28564 | 64772 | 28848 |
| BAC | 33445 | 4937 | 14113 | 12582 | 36741 | 12556 |
| BS | 23934 | 12955 | 14084 | 23102 | 41724 | 22090 |
| DORsm | 18224 | 10397 | 9458 | 12750 | 33125 | 15360 |
| AD | 32506 | 15030 | 15181 | 12315 | 32209 | 11114 |
| ARH | 27038 | 13899 | 10118 | 14098 | 39273 | 21029 |
| ADP | 46301 | 24031 | 22182 | 14057 | 41098 | 12577 |
| AHN | 23459 | 11668 | 9114 | 12842 | 19713 | 21558 |
| A13 | 38540 | 53371 | 18192 | 38781 | 102410 | 40203 |
| MB | 42072 | 16804 | 19880 | 27295 | 49630 | 28389 |
| IC | 39670 | 17201 | 21194 | 19085 | 51554 | 17441 |
| APN | 22274 | 19255 | 14006 | 8470 | 25487 | 7117 |
| SNc | 36840 | 9585 | 20578 | 11117 | 35509 | 9367 |
| KF | 43678 | 7455 | 17965 | 14252 | 52332 | 12439 |
| Acs5 | 36638 | 13273 | 19135 | 15717 | 21821 | 14029 |
| CS | 54113 | 14353 | 25220 | 13968 | 24724 | 12519 |
| AP | 36638 | 13273 | 19135 | 15717 | 21821 | 14029 |
| ACVI | 54113 | 14353 | 25220 | 13968 | 24724 | 12519 |

**Supporting table 4:**
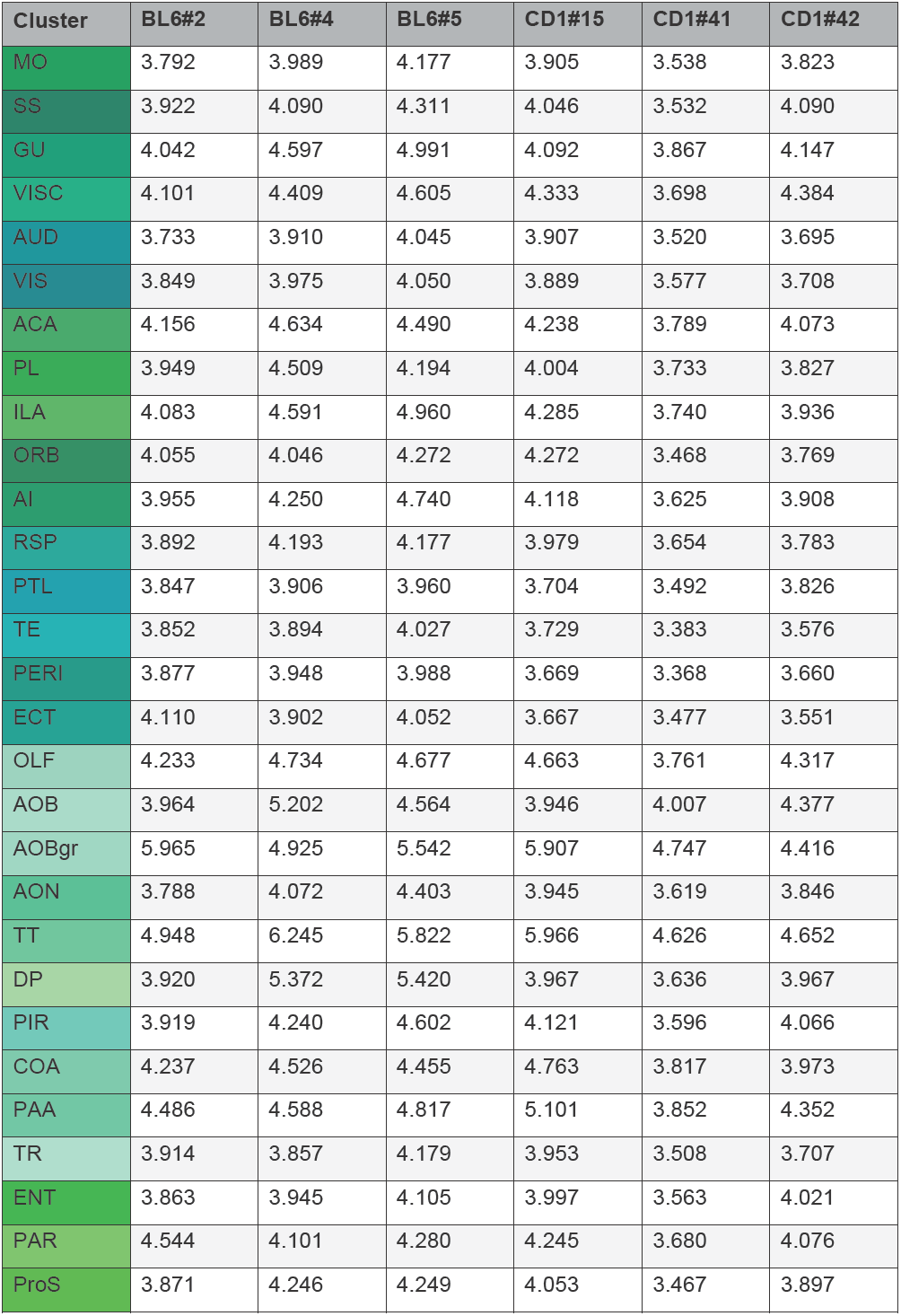
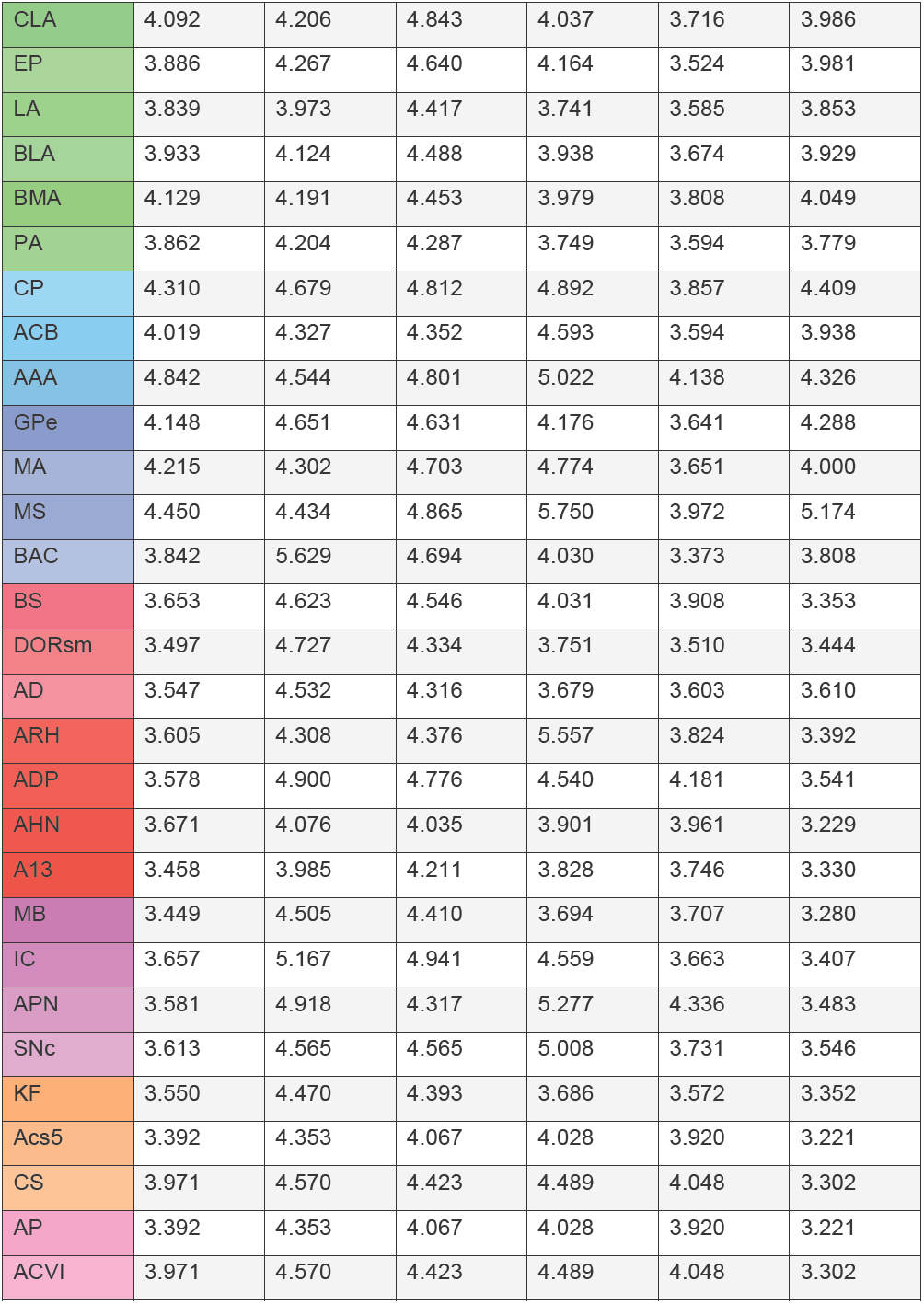
Quantification of the radii in the cleared C57BL/6J and CD1-E samples, units are *μm*.

| Cluster | BL6#2 | BL6#4 | BL6#5 | CD1#15 | CD1#41 | CD1#42 |
| --- | --- | --- | --- | --- | --- | --- |
| MO | 3.792 | 3.989 | 4.177 | 3.905 | 3.538 | 3.823 |
| SS | 3.922 | 4.090 | 4.311 | 4.046 | 3.532 | 4.090 |
| GU | 4.042 | 4.597 | 4.991 | 4.092 | 3.867 | 4.147 |
| VISC | 4.101 | 4.409 | 4.605 | 4.333 | 3.698 | 4.384 |
| AUD | 3.733 | 3.910 | 4.045 | 3.907 | 3.520 | 3.695 |
| VIS | 3.849 | 3.975 | 4.050 | 3.889 | 3.577 | 3.708 |
| ACA | 4.156 | 4.634 | 4.490 | 4.238 | 3.789 | 4.073 |
| PL | 3.949 | 4.509 | 4.194 | 4.004 | 3.733 | 3.827 |
| ILA | 4.083 | 4.591 | 4.960 | 4.285 | 3.740 | 3.936 |
| ORB | 4.055 | 4.046 | 4.272 | 4.272 | 3.468 | 3.769 |
| AI | 3.955 | 4.250 | 4.740 | 4.118 | 3.625 | 3.908 |
| RSP | 3.892 | 4.193 | 4.177 | 3.979 | 3.654 | 3.783 |
| PTL | 3.847 | 3.906 | 3.960 | 3.704 | 3.492 | 3.826 |
| TE | 3.852 | 3.894 | 4.027 | 3.729 | 3.383 | 3.576 |
| PERI | 3.877 | 3.948 | 3.988 | 3.669 | 3.368 | 3.660 |
| ECT | 4.110 | 3.902 | 4.052 | 3.667 | 3.477 | 3.551 |
| OLF | 4.233 | 4.734 | 4.677 | 4.663 | 3.761 | 4.317 |
| AOB | 3.964 | 5.202 | 4.564 | 3.946 | 4.007 | 4.377 |
| AOBgr | 5.965 | 4.925 | 5.542 | 5.907 | 4.747 | 4.416 |
| AON | 3.788 | 4.072 | 4.403 | 3.945 | 3.619 | 3.846 |
| TT | 4.948 | 6.245 | 5.822 | 5.966 | 4.626 | 4.652 |
| DP | 3.920 | 5.372 | 5.420 | 3.967 | 3.636 | 3.967 |
| PIR | 3.919 | 4.240 | 4.602 | 4.121 | 3.596 | 4.066 |
| COA | 4.237 | 4.526 | 4.455 | 4.763 | 3.817 | 3.973 |
| PAA | 4.486 | 4.588 | 4.817 | 5.101 | 3.852 | 4.352 |
| TR | 3.914 | 3.857 | 4.179 | 3.953 | 3.508 | 3.707 |
| ENT | 3.863 | 3.945 | 4.105 | 3.997 | 3.563 | 4.021 |
| PAR | 4.544 | 4.101 | 4.280 | 4.245 | 3.680 | 4.076 |
| ProS | 3.871 | 4.246 | 4.249 | 4.053 | 3.467 | 3.897 |

**Supporting table 4:** Quantification of the radii in the cleared C57BL/6J and CD1-E samples, units are $\mu m$ .
|  |  |  |  |  |  |  |
| --- | --- | --- | --- | --- | --- | --- |
| CLA | 4.092 | 4.206 | 4.843 | 4.037 | 3.716 | 3.986 |
| EP | 3.886 | 4.267 | 4.640 | 4.164 | 3.524 | 3.981 |
| LA | 3.839 | 3.973 | 4.417 | 3.741 | 3.585 | 3.853 |
| BLA | 3.933 | 4.124 | 4.488 | 3.938 | 3.674 | 3.929 |
| BMA | 4.129 | 4.191 | 4.453 | 3.979 | 3.808 | 4.049 |
| PA | 3.862 | 4.204 | 4.287 | 3.749 | 3.594 | 3.779 |
| CP | 4.310 | 4.679 | 4.812 | 4.892 | 3.857 | 4.409 |
| ACB | 4.019 | 4.327 | 4.352 | 4.593 | 3.594 | 3.938 |
| AAA | 4.842 | 4.544 | 4.801 | 5.022 | 4.138 | 4.326 |
| GPe | 4.148 | 4.651 | 4.631 | 4.176 | 3.641 | 4.288 |
| MA | 4.215 | 4.302 | 4.703 | 4.774 | 3.651 | 4.000 |
| MS | 4.450 | 4.434 | 4.865 | 5.750 | 3.972 | 5.174 |
| BAC | 3.842 | 5.629 | 4.694 | 4.030 | 3.373 | 3.808 |
| BS | 3.653 | 4.623 | 4.546 | 4.031 | 3.908 | 3.353 |
| DORsm | 3.497 | 4.727 | 4.334 | 3.751 | 3.510 | 3.444 |
| AD | 3.547 | 4.532 | 4.316 | 3.679 | 3.603 | 3.610 |
| ARH | 3.605 | 4.308 | 4.376 | 5.557 | 3.824 | 3.392 |
| ADP | 3.578 | 4.900 | 4.776 | 4.540 | 4.181 | 3.541 |
| AHN | 3.671 | 4.076 | 4.035 | 3.901 | 3.961 | 3.229 |
| A13 | 3.458 | 3.985 | 4.211 | 3.828 | 3.746 | 3.330 |
| MB | 3.449 | 4.505 | 4.410 | 3.694 | 3.707 | 3.280 |
| IC | 3.657 | 5.167 | 4.941 | 4.559 | 3.663 | 3.407 |
| APN | 3.581 | 4.918 | 4.317 | 5.277 | 4.336 | 3.483 |
| SNc | 3.613 | 4.565 | 4.565 | 5.008 | 3.731 | 3.546 |
| KF | 3.550 | 4.470 | 4.393 | 3.686 | 3.572 | 3.352 |
| Acs5 | 3.392 | 4.353 | 4.067 | 4.028 | 3.920 | 3.221 |
| CS | 3.971 | 4.570 | 4.423 | 4.489 | 4.048 | 3.302 |
| AP | 3.392 | 4.353 | 4.067 | 4.028 | 3.920 | 3.221 |
| ACVI | 3.971 | 4.570 | 4.423 | 4.489 | 4.048 | 3.302 |

